# Chronic Stress Increases Adiposity and Anxiety in Rats with Decreased Expression of *Krtcap3*

**DOI:** 10.1101/2023.06.24.546378

**Authors:** Alexandria Szalanczy, Mackenzie Fitzpatrick, Angela Beeson, Trangdai Bui, Christina Dyson, Seth Eller, Julia Landry, Christina Scott, Michael Grzybowski, Jason Klotz, Aron M Geurts, Jeff L Weiner, Eva E Redei, Leah C Solberg Woods

## Abstract

We previously identified *Keratinocyte-associated protein 3*, *Krtcap3*, as a novel adiposity gene but subsequently found that its impact on adiposity may depend on environmental stress. To more thoroughly understand the connection between *Krtcap3*, adiposity, and stress, we exposed wild-type (WT) and *Krtcap3* knock-out (KO) rats to chronic stress then measured adiposity and behavioral outcomes. We found that KO rats displayed lower basal stress than WT rats under control conditions and exhibited the expected responses to chronic stress exposure. Specifically, stress-exposed KO rats gained more weight, consumed more food when socially isolated, and displayed more anxiety-like behaviors relative to control KO rats. Meanwhile, there were minimal differences between control and stressed WT rats. At study conclusion stress-exposed KO rats had increased corticosterone (CORT) relative to control KO rats with no differences between WT rats. In addition, KO rats, independent of prior stress exposure, had an increased CORT response to removal of their cage-mate (psychosocial stress), which was only seen in WT rats when exposed to chronic stress. Finally, we found differences in expression of the glucocorticoid receptor, *Nr3c1*, in the pituitary and colon between control and stress-exposed KO rats that were not present in WT rats. These data support that *Krtcap3* expression affects stress response, potentially via interactions with *Nr3c1*, with downstream effects on adiposity and behavior. Future work is necessary to more thoroughly understand the role of *Krtcap3* in the stress response.

## Introduction

Obesity continues to be a national and global health crisis. Data from 2021 show that over 40% of adults are characterized as having obesity (Stierman 2021), which is commonly attributed to decreases in diet quality and increases in sedentary lifestyles. Obesity is a complex disease that results from interaction of genetic and environmental factors. The genetic architecture of obesity remains poorly understood, despite the hundreds of genetic loci that have been identified (Abadi, Alyass et al. 2017, Yengo, Sidorenko et al. 2018, Pulit, Stoneman et al. 2019, Rohde, Keller et al. 2019). Importantly, environmental factors include more than diet and exercise but are not as frequently investigated. For example, stress can have profound effects on eating behavior, fat deposition, and mental health, which may induce or propagate the well-established vicious cycle between obesity, dysregulated metabolic health, and altered mental health (Lee, Cardel et al. 2000, Torres and Nowson 2007, Calabrese, Molteni et al. 2009, Peckett, Wright et al. 2011, Scott, Melhorn et al. 2012, van der Valk, Savas et al. 2018, Lahdepuro, Savolainen et al. 2019, Kahan 2020, Xiao, Liu et al. 2020, Wang, Cheng et al. 2022). Though the connection between obesity and stress is well-known, only a small amount of research has specifically investigated the genetic connections between obesity and stress. Prior work has shown that there is a genetic component to the stress response (Ising and Holsboer 2006, Solberg, Baum et al. 2006, van den Bos, Harteveld et al. 2009, Henckens, Klumpers et al. 2016, Flati, Gioiosa et al. 2020), which raises the possibility that genetic variation in stress reactivity may connect to variation in adiposity.

We first identified *Keratinocyte-associated protein 3* (*Krtcap3*) as a candidate gene for adiposity using a genome-wide association study (GWAS) in rats (Keele, Prokop et al. 2018, Chitre, Polesskaya et al. 2020). Initial work in an *in vivo Krtcap3* knock-out (KO) model supported these GWAS findings, demonstrating that female KO rats had increased food intake and adiposity compared to wild-type (WT) controls. However, we were unable to replicate these initial results in a subsequent study, which may have been due to changes in environmental stress (Szalanczy, Giorgio et al. 2023). COVID-19 shut-downs caused large environmental differences in rat housing conditions between the two studies, including the completion of a nearby construction project, that may have decreased the stress of the rats. We found that WT rats ate more, had increased adiposity, and had decreased serum corticosterone (CORT) in the second study, supporting a decrease in stress. The same phenotypes were not altered in KO rats between the two studies. We proposed that *Krtcap3* may play a role in stress response with secondary effects on adiposity (Szalanczy, Giorgio et al. 2023).

In the current study, we sought to investigate the relationship between *Krtcap3* expression, adiposity, and stress. We expected that WT rats would show a reduction in food intake and adiposity when exposed to chronic low-grade physical stress, while KO rats would remain resistant to the effects of chronic stress. When naïve to stress, we anticipated that WT and KO rats would have similar adiposity measures. We also included several behavioral tests as chronic stress has been shown to worsen mental health in humans (Calabrese, Molteni et al. 2009, Lahdepuro, Savolainen et al. 2019) and alter emotion-like behavior in rodents (Sequeira-Cordero, Salas-Bastos et al. 2019, Atrooz, Alkadhi et al. 2021). We hypothesized that WT rats would show worsened behavioral outcomes in response to chronic stress relative to KO rats. We initially sought to induce stress by mimicking the increased noise conditions from the first study by exposing rats to experimentally controlled, loud auditory stimuli. When we did not see increases in serum corticosterone or changes in adiposity, we began a second arm of stress delivery using a modified unpredictable chronic mild stress (UCMS) paradigm (Willner 2017) where rats were exposed to a variety of mild stressors six days of the week.

Contrary to expectations, we found that stress more strongly increased adiposity and heightened anxiety-like behaviors in KO rats, not WT. We suspect that use of the stress-sensitive Wistar-Kyoto (WKY) strain (Redei, Udell et al. 2022) may be significant in interpreting these results. While control rats were not exposed to the two stress protocols, they participated in numerous metabolic and behavior tests that may have induced stress. WKY rats may not have been able to distinguish between the chronic stress protocol (stress-exposed) versus the stress of the study design itself (control rats), while KO rats were better able to distinguish between types of stress. While we were ultimately unable to replicate the previous adiposity findings, this study demonstrates that WT and *Krtcap3*-KO rats have different metabolic and behavioral responses to stress, supporting a role of *Krtcap3* in the stress response.

## Methods

### Animals

We previously generated a whole-body *in vivo Krtcap3*-KO on the WKY (WKY/NCrl; RGD_1358112) inbred rat strain (WKY-Krtcap3^em3Mcwi^) and established a breeding colony at Wake Forest University School of Medicine (WFUSOM) in 2019 (Szalanczy, Goff et al. 2022). At Building A of WFUSOM, rats were housed in ventilated cages (46 cm x 24 cm x 20 cm) at 22°C in a 12 h light/dark cycle (dark from 18:00 to 6:00) at standard temperature and humidity conditions, and given *ad libitum* access to water. Bedding was 0.25 in corn cob with a paper puck for enrichment. Breeder rats and experimental rats prior to high-fat diet (HFD) start were given *ad libitum* standard chow diet (Lab Diet, Prolab RMH 3000, Catalog #5P00). At six weeks of age, experimental WT and KO rats were placed on experimental diet as described below (n = 8 female rats per genotype and stress condition).

Experimental animals in the stress study were housed at Building A in the same room as the breeding colony up until approximately 23 weeks of age, at which point they were transferred to the nearby Building B, which is better set up for rodent behavioral testing. At Building B, rats were housed in a male/female room in non-ventilated, open-air wire top cages (26.5 cm x 48 cm x 20 cm) with aspen shavings. Rats were maintained on the same 12 h light/dark cycle (dark from 18:00 to 6:00) and access to food and water remained *ad libitum*. Building B is an older building that does not allow temperature and humidity to be as tightly controlled as Building A, but conditions are appropriately managed with animal care health observation and veterinary oversight.

We also sought to evaluate *Krtcap3* expression in multiple tissues between adolescent WT and KO rats from the colony without diet or stress exposure. Female WT and KO (n = 4 per genotype) rats were weaned at three weeks of age and placed into mixed-genotype cages with littermates, with two to four rats per cage. Rats were housed in Building A and maintained on the same chow diet as breeders. One week later, rats were euthanized, described below.

All experiments were performed using a protocol approved by the Institutional Animal Care and Use Committee at WFUSOM.

### Genotyping

Experimental rats were genotyped at the Medical College of Wisconsin (MCW) as described elsewhere (Szalanczy, Giorgio et al. 2023).

### Stress Study Design

Female experimental rats were weaned at three weeks of age, weighed, and placed two per cage in same-sex, same-genotype cages. Rats were randomly assigned to either a control or stress group, as described in detail below. Stress study rats were initially maintained on the same diet as breeders (see above), and body weight was recorded weekly starting at four weeks of age. To remain consistent with our previous studies (Szalanczy, Goff et al. 2022, Szalanczy, Giorgio et al. 2023), all rats began a HFD (60% kcal fat; ResearchDiet D12492) at six weeks of age. Rats were allowed access to diet *ad libitum* with cage-wide food intake recorded weekly.

From the time of weaning, the study lasted for 22 weeks (**Figure 1**). For rats assigned into the stress group, a mild stress exposure began at 22 days of age and lasted for approximately 12 weeks. Control rats underwent the same metabolic and behavioral phenotyping as stress rats, but were not exposed to the stress protocols. The first EchoMRI analysis was conducted three weeks after stress onset, at HFD start, while an acute restraint test took place ten weeks after stress onset. The first two behavioral tests were the novelty suppressed feeding test (NSF) conducted 11.5 weeks after stress onset and the forced swim test (FST) conducted a week later. Another EchoMRI analysis was run 13 weeks after stress onset, leading immediately into a week-long period of individual housing to assess food intake. Following return to cage-mate, UCMS began 15 weeks after initial stress onset, and continued for four weeks, concluding with another EchoMRI analysis and an intraperitoneal glucose tolerance test (IPGTT). After rats were transferred to Building B they were given an open field test (OFT) 20.5 weeks after initial stress onset. 22 weeks after stress onset rats were euthanized (Sac). Blood was collected at three points to measure CORT responses: at the acute restraint test, immediately prior to UCMS, and at study termination.

**Figure 1.**
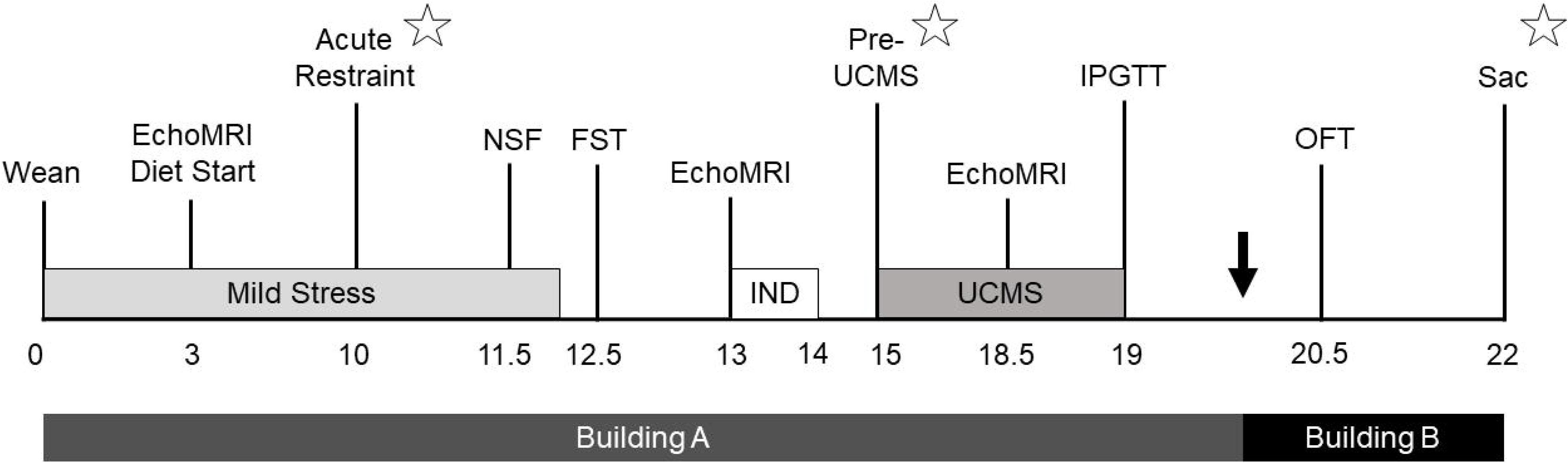
Study timeline. Timeline outlining study design, with weeks relative to stress start shown. Rats were housed in Building A from birth to approximately 20 weeks from stress start, and then Building B until study completion. Mild stress was administered starting after weaning and continued for 12 weeks, while four weeks of unpredictable chronic mild stress (UCMS) started 15 weeks after initial stress exposure. Metabolic phenotyping included EchoMRI analysis (EchoMRI), a week of individual housing (IND) to measure individual food intake, an intraperitoneal glucose tolerance test (IPGTT), and euthanasia (Sac). Behavioral tests included the novelty suppressed feeding test (NSF), forced swim test (FST), and open field test (OFT). Corticosterone was assessed 10 weeks after stress start at an acute restraint test, prior to UCMS 15 weeks into the study, and then at euthanasia (represented by star symbols).

### Stress Protocol Designs

#### Mild Stress Protocol

We first started with a mild stress protocol that emphasized noise exposure in an effort to mimic the increased noise in the vivarium prior to COVID-19 shut-downs (Szalanczy, Goff et al. 2022). White noise exposure has previously been shown to increase CORT in rodents (Burow, Day et al. 2005, Samson, Sheeladevi et al. 2007). We modified other stress-delivery protocols that exposed rodents to white noise (Campeau, Dolan et al. 2002, Burow, Day et al. 2005, Samson, Sheeladevi et al. 2007) by placing rats in a small, well-lit, enclosed room, 12 in away from a noise machine (Serene Evolution White Noise Machine, Amazon ASIN B08FZSGFSK) set to max volume (70 dB) that was low enough to avoid hearing loss (Escabi, Frye et al. 2019). In order to minimize acclimatization (Masini, Day et al. 2008), the noise was cycled through different types of noise, different intervals, and different times of the day. The types of noise included white noise, brown noise, thunder, vacuum, and crowd. Noise exposure lasted from one to three hours, delivered anytime between 8:00 to 17:00, and was randomly broken into the following intervals: straight, one hour on and 30 min off, 30 min on and 30 min off, 40 min on and 20 min off, 15 min on and 15 min off. The noise machine could be turned on and off from outside the room.

In our first *in vivo* study conducted from 2019-2020, KO rats were larger than WT rats by six weeks of age (Szalanczy, Goff et al. 2022). In the current study, we did not see the anticipated changes to body weight at six weeks of age (HFD start) and concluded that the noise procedure described above was too mild to causes increases in CORT. We then altered the stress design by decreasing exposure to white noise to two days a week but adding exposure to an additional stressor one time per week, modified from those used in the UCMS paradigm (Willner 2017): 30 min of restraint, a five min swim in 22 °C water, a cage tilted at 45° for two hours, five min exposure to a cool air stream from a hair dryer, or eight hours in a flooded cage (500 mL water added to bedding). This mild stress protocol lasted for nine weeks after diet start, through the first round of behavioral tests discussed below.

#### Unpredictable Chronic Mild Stress (UCMS)

UCMS is an established, validated protocol for generating stress in rodents, inducing behavioral changes that correlate with increased anxiety-like and depressive-like symptoms (Willner 2017, Willner 2017). The UCMS procedure that we used was taken from the literature (Isingrini, Camus et al. 2010, Frisbee, Brooks et al. 2015, Monteiro, Roque et al. 2015, Willner 2017, Burstein and Doron 2018, Alqurashi, Hindi et al. 2022), with modifications: rats remained pair-housed during the duration of stress delivery and stressors were administered six days per week instead of seven. Stressors were administered in either the housing room or different procedure rooms and included: a two-hour restraint in a flat-bottomed tube (see below), 15-min exposure to a cotton ball soaked in fox urine, a five min swim in water approximately 15 °C, overnight exposure to soiled breeder bedding, five min exposure to a cool air stream from a hair dryer, three to eight hours in a flooded cage, and two to three hours in a cage tilted at approximately 45°. UCMS began on Monday and ended just over four weeks later with an overnight fast for a glucose tolerance test (discussed below). The exact schedule of procedures for UCMS is given in **Table I**. As with other UCMS protocols (Isingrini, Camus et al. 2010, Frisbee, Brooks et al. 2015, Monteiro, Roque et al. 2015, Willner 2017, Alqurashi, Hindi et al. 2022) rats did not see the same stressor two days in a row, and timing of stress delivery varied from day to day. With the exception of overnight exposure to soiled bedding, all other procedures were given from 8:00 to 17:00. Four researchers contributed to UCMS stress delivery.

**Table I.** Sequence of unpredictable chronic mild stress (UCMS) procedures. UCMS stressors included a two-hour restraint in a plastic tube (Restraint), a 15-minute exposure to fox urine (Fox), a five-minute swim in cool (15 °C) water (Cool Swim), overnight exposure to soiled bedding from breeder cages (Soiled), five-minute exposure to a cool air stream from a hair dryer (Dryer), three to eight hours exposed to a flooded cage (Flood), and two to three hours exposed to a cage tilt at 45° (Tilt). UCMS began on a Monday, and stressors were delivered Sunday through Monday with Saturdays off. Stressors, excepting overnight exposure to soiled bedding, were administered randomly from 8:00 to 17:00, and rats did not see the same stressors twice in a row. On the last Sunday of the UCMS period rats had a third EchoMRI analysis, and UCMS concluded with an overnight fast and an intraperitoneal glucose tolerance test (IPGTT).

| Week | Sun | Mon | Tues | Wed | Thurs | Fri | Sat |
| --- | --- | --- | --- | --- | --- | --- | --- |
| 1 |  | Restraint | Fox | Cool Swim | Soiled | Dryer |  |
| 2 | Flood | Soiled | Cool Swim | Tilt | Flood | Fox |  |
| 3 | Cool Swim | Dryer | Restraint | Soiled | Tilt | Flood |  |
| 4 | Soiled | Cool Swim | Fox | Flood | Dryer | Restraint |  |
| 5 | EchoMRI | Dryer | Fast | IPGTT |  |  |  |

### Metabolic Phenotyping

Throughout the course of the study, body weight, and cage-wide food intake were measured weekly. EchoMRI analysis (EchoMRI LLC, Houston, TX) was conducted at three, 13, and 18.5 weeks after first stress exposure (Szalanczy, Goff et al. 2022). Immediately following the EchoMRI analysis 13 weeks after stress onset, rats began a seven day-long period of individual housing to measure food intake as previously described (Szalanczy, Giorgio et al. 2023). 19 weeks after stress onset, rats were fasted overnight before being administered an intraperitoneal glucose tolerance test (IPGTT) as performed elsewhere (Szalanczy, Goff et al. 2022, Szalanczy, Giorgio et al. 2023). We measured blood glucose (Contour Next EZ) at fasting and 15, 30, 60, 90, and 120 minutes after a 1 mg/kg glucose injection and calculated glucose area-under-the-curve (AUC) (Szalanczy, Goff et al. 2022, Szalanczy, Giorgio et al. 2023).

### Blood Collection for measurement of basal and restraint CORT

#### Acute Restraint Test

To determine if there were differences in basal CORT due to the mild stress protocol, we collected blood 10 weeks after mild stress onset. Rats were also subjected to a 30 min acute restraint test to investigate if there were different CORT responses by genotype to an acute physical stressor. The test was performed in the housing room in the morning (9:00-10:00) with two researchers. Rats were removed from the cage at the same time, placed in either a plastic flat-bottomed restraint tube (Braintree Scientific Cat #FB-ML or Cat #FB-L) or a restraint bag (Fisher Scientific Cat #14370112), and their tails nicked to collect serum (Sarstedt Inc MicrovetteCB 300 LH). Basal collection was completed within 90 sec of the rats being removed from the cage. After completing the basal collection, tails were taped to the surface for further restraint. 10 min later, rats were again bled from the tail, still restrained. After 30 min total, rats were bled a final time then removed from the restraint and returned to the home cage. Two cages could be run in the same 30 min chunk, staggered 5 min apart.

#### Pre-UCMS

We collected blood approximately 15 weeks after stress onset to obtain a basal CORT measurement prior to starting UCMS. Starting in the morning (9:00-10:00), rats were briefly removed from their cage, their tail nicked, and serum collected. This was performed in the housing room with two researchers—both rats were removed from the cage at the same time, and blood collection was completed within 90 s of removal. Rats were then either returned to the home cage (control rats) or remained restrained for another 90 min as the first stressor of UCMS (stressed rats).

#### Post-UCMS

We collected blood for basal CORT at euthanasia, described below under Tissue Harvest.

### Behavioral Analyses

#### Novelty Suppressed Feeding Test (NSF)

About 11 weeks after mild stress exposure began, rats were administered the NSF to measure anxiety-like behaviors such as hyponeophagia (Dulawa and Hen 2005) and exploratory behaviors. Rats were fasted for 24 h before the test began, then brought to a procedure room to acclimate for 30 min before the test started. A large box (67 cm x 61 cm x 46 cm) was placed in the center of the room under an overhead light with a camera positioned at one edge to look down over the box. The sides of the box were covered in black construction paper to minimize outside distractions. The bottom of the box was marked in red tape with two concentric circles (diameters 16 and 47 cm) and six lines radiating at 60° from the edge of the inner circle, dividing the box into 13 sections (McAuley, Stewart et al. 2009).

The box was cleaned with 70% EtOH between rats. A small dish with three pellets of HFD was positioned in the center of the box and taped down to prevent movement. The video camera was started, a rat was removed from its cage and placed in the left corner facing the walls of the box, and then the researcher left the room and started a timer. The rat was left alone in the box for 15 min, at which point the researcher returned to the room to return the rat to its home cage. The box was cleaned with 70% ethanol between rats.

Researchers scoring these measures were blinded to the genotype and stress exposure of the rats, and the same researcher scored the same phenotype across all rats. Latency to feed, or how long it took the rat to begin eating, was defined as the rat chewing for at least four consecutive seconds. From that point, total time spent feeding was measured based on the total time the rat spent chewing. The number of line crossings, the number of center approaches, and time spent in the center were also measured. A line crossing was defined as all four paws crossing from one block into another. The smaller circle was considered the center of the box, and due to the blockage of the food dish, a rat was considered to have entered the center when both front paws were on or over the line. Time in center began when two front paws were in the center, and ended when at least one paw left the center. Eating within the center was also counted within the center time. Increased latency to feed, decreased total feeding time, fewer line crossings, and decreased center time are all considered anxiety-like behaviors in rodents (Dulawa and Hen 2005, Seibenhener and Wooten 2015).

#### Forced Swim Test (FST)

Three days after the NSF, rats participated in a FST to measure passive coping to stress (Commons, Cholanians et al. 2017) and depressive-like behavior (Redei, Udell et al. 2022). Rats were placed in a tank of water (25 °C ± 2, diameter 28.8 cm, height 49.8 cm, water depth 39 cm) for 15 min on Day 1 and 5 min on Day 2 as previously described (Solberg, Baum et al. 2004). Video recording of the first five minutes of Day 1 and the full five minutes of Day 2 were used to manually score movements made by the rat at 5 s intervals: immobility (floating or bracing) and mobility (swimming, climbing, or diving). Increased immobility is associated with increased passive coping (Commons, Cholanians et al. 2017) and/or depressive-like behavior (Redei, Udell et al. 2022).

#### Open Field Test (OFT)

20.5 weeks after stress exposure began, and 1.5 weeks after the conclusion of UCMS, an OFT was administered to measure locomotor activity and anxiety-like behaviors in the rats driven by the competing urges of exploring new environments but avoiding open, well-lit spaces (Seibenhener and Wooten 2015). The test was conducted in Plexiglas chambers (41.5 cm x 41.5 cm x 30 cm). The center of the box was measured as a 20.5 cm x 20.5 cm square. At the start of the test, rats were placed in the chambers equipped with Omnitech Superflex Sensors (Omnitech Electronics, Inc., Columbus, OH), which utilize arrays on infrared photodetectors located at regular intervals along each way of the chambers. The chamber walls are solid and contained within sound-attenuating boxes with a 7.5-W white light to illuminate the arena. Exploratory activity in this environment was measured for 30 min and the data analyzed in five-minute time bins. The following activities were recorded: total distance moved, total time spent moving, number of rears and time spent rearing, number of center approaches, time spent in the center of the box, and distance traveled in the center of the box.

### Tissue Harvest

#### Stress Study

After 22 weeks of stress exposure and beginning at 8:00, rats were euthanized via decapitation after a 4 h fast. Rats were transferred to the anteroom of the necropsy suite 15 min prior to the start of the euthanasia protocol, and were euthanized one at a time. Weight gain was calculated as the difference between the final body weight of the rats following the fast and the body weight at study start. Plasma was collected from trunk blood (Fisher Scientific Cat #02-657-32) and saved at -80 C. Body length from nose to anus and tail length from anus to tail tip were measured with a ruler. The brain, retroperitoneal (RetroFat), and liver were dissected, weighed, and snap-frozen. The pituitary and adrenal glands as well as the ovaries and sections of the ileum and colon were dissected and snap-frozen without being weighed. Adrenal glands were later weighed on a more precise scale. Kidneys, parametrial fat (ParaFat), and omental/mesenteric fat (OmenFat) were weighed but not saved.

#### Adolescent rats for *Krtcap3* expression

To confirm the *Krtcap3*-KO in multiple tissues, four week-old control rats were transferred to the anteroom of the necropsy suite 30 min prior to euthanasia, without fasting. Rats were euthanized with CO_2_ for five min, then decapitated. The brain, RetroFat, liver, pituitary gland, adrenal gland, ovaries, a section of the ileum, and a section of the colon were dissected and immediately snap-frozen in liquid nitrogen.

### Adrenocorticotropic Hormone & Corticosterone

To measure adrenocorticotropic hormone (ACTH) in plasma collected at euthanasia, we used a sandwich ELISA kit (AbCam Ref #ab263880). Plasma samples were diluted 1:4 in 1X Cell Extraction Buffer PTR. The plate was incubated for 15 min in the TMB substrate, then Stop Solution was added, and the plate was analyzed at 450 nm against a 4-parameter standard curve.

We used a CORT competitive ELISA kit (ThermoFisher Ref # EIACORT) to analyze serum/plasma corticosterone collected throughout the study. As guided by the manufacturer’s instructions, samples were diluted at least 1:100, and analyzed at 450 nm against a 4-parameter standard curve.

### RNA Extraction

RNA was extracted from fatty tissues using the RNeasy Lipid Tissue Mini Kit (Qiagen Cat # 74804). Non-fatty tissue, such as liver or intestine, were extracted by Trizol.

### Real Time Quantitative PCR

*Pro-opiomelanocortin* (*Pomc*) expression from the pituitary was measured between rats from the stress study to determine if there were genotype or stress exposure-driven differences on expression of the precursor to ACTH. We also examined expression of the glucocorticoid receptor (GR) *nuclear receptor subfamily 3 group C member 1* (*Nr3c1*) in liver, colon, and pituitary between females of the main study to assess changes in CORT receptor expression by genotype and stress exposure. Colon and pituitary have high *Krtcap3* expression (**Supplementary Figure 1**) while we previously investigated *Nr3c1* expression in the liver (Szalanczy, Giorgio et al. 2023). We also investigated expression of *hydroxysteroid 11-beta dehydrogenase 2* (*Hsd11*β*2*), responsible for catalyzing CORT deactivation, in the colon to determine if there were differences in CORT processing.

We then measured *Krtcap3* expression between control and stress-exposed WT females in pituitary, adrenal, and colon to determine if expression had changed following exposure to the stress protocols. Results from the adolescent rats (described below) confirmed KO rats did not have *Krtcap3* expression in these tissues, and they were excluded from this analysis.

We assessed *Krtcap3* expression in adolescent females to confirm that *Krtcap3* expression was significantly knocked-out in multiple tissues. We examined expression in tissues pertinent to metabolism or stress response: pituitary, adrenal, liver, whole hypothalamus, ileum, ovaries, and RetroFat.

To measure gene expression, either *Gapdh* or β*-actin* were used as housekeeping genes. Primers for all genes are found in **Table II**. Fold change of the corresponding gene of interest transcript was calculated by the following equation:

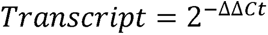

**Table II.**
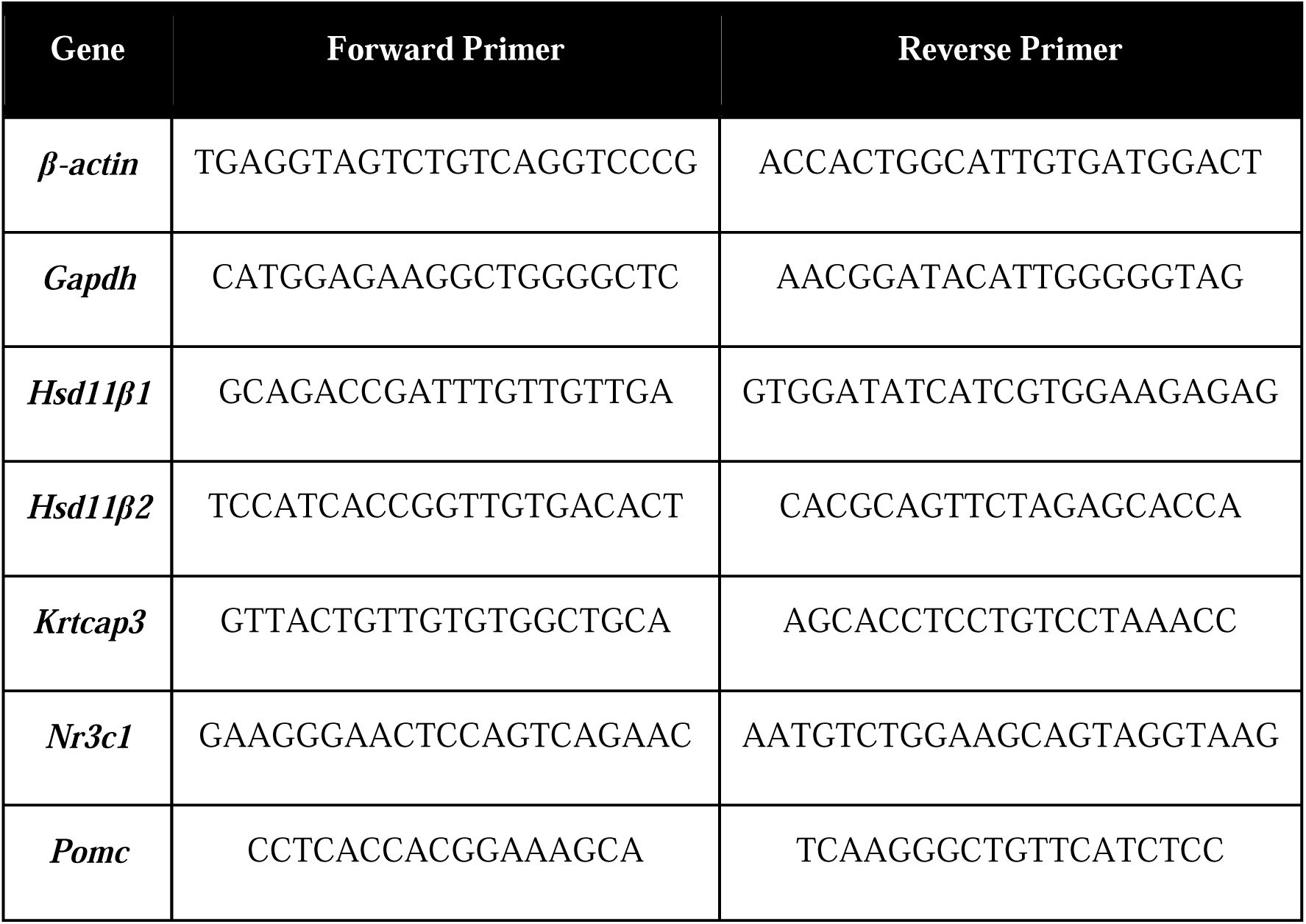
Primer sequences. 5’ ➔ 3’

Where ΔCt was the difference between the crossing threshold (Ct) of the gene of interest and the housekeeping gene, and ΔΔCt the difference between each sample ΔCt and the average control ΔCt. For analysis of the adolescent rats, the average control ΔCt was the mean ΔCt of WT rats. In gene expression analyses in the main study, the average control ΔCt was the mean ΔCt of control WT rats.

### Statistical Analysis

All data were analyzed in R (1.4.1103). Outliers were assessed either by Grubb’s test or by 1.5 * the interquartile range for phenotypes where n < 6 and were removed. Distribution was assessed by the Shapiro-Wilks test and data were transformed to reflect a normal distribution if necessary. Homogeneity of variance was assessed by Levene’s test.

For adiposity, behavioral, and gene expression results, data were first analyzed considering all four groups together. If ANOVA analysis indicated relevant interactions, the data were deconstructed appropriately. However, several phenotypes showed a visually stronger effect in one genotype relative to the other, even if ANOVA could not detect an interaction. In these cases, according to our *a priori* hypothesis that WT and KO rats would respond differently to stress, we split the data by genotype and assessed effect of stress.

Single point adiposity and behavioral measurements were first assessed by a two-way ANOVA, where one factor was genotype (WT v KO) and the other was stress exposure (control v stress). Growth curves were analyzed by a two-way repeated measures ANOVA, where the effect of genotype and stress exposure were examined over time. HFD consumption during the mild stress protocol (weeks 3-12 of study) was calculated as the food consumed by each cage per day for each week and then averaged. Food consumed during the week of individual housing was calculated as the average food consumed per day and the total food consumed during the week for each rat. Food consumed during the month of UCMS was calculated as the food consumed per cage per day each week and then averaged as well as the total food consumed per cage for the month. If there was an interaction, or visually WT and KO rats responded differently, we split the data by genotype to evaluate the effect of stress or the effect of stress and time together.

Based on our previous study (Szalanczy, Giorgio et al. 2023) we anticipated there being three factors to consider when analyzing CORT and ACTH data from euthanasia: genotype, stress exposure, and euthanasia order (if the rat was euthanized first or second within a cage). We first examined CORT and ACTH from euthanasia using a two-way ANOVA, with only data from Rat 1 which we presumed to represent the basal condition, removing the consideration for the factor of order. The data were then analyzed by a three-way ANOVA and split according to significant interactions. We did not include rat order as a factor in analyses of CORT collected from the acute restraint test or prior to the UCMS protocol as both rats were handled at the same time. We used a mixed effects model to analyze CORT during the acute restraint, with the factors of genotype, stress exposure, and time restrained; Tukey’s multiple comparisons test compared CORT between the time points. Given the large effect of time, we then also assessed CORT at each time point separately with a two-way ANOVA. To evaluate CORT prior to UCMS exposure, we used a two-way ANOVA.

In the NSF, latency to feed and total time spent feeding could not be analyzed by a two-way ANOVA because there was a significant ceiling effect, as over one-quarter of the rats did not participate in the test in the given timeframe. Instead, we chose to re-conceptualize the latency to feed and time spent feeding phenotypes as participation in the test, and to then analyze by methods similar to those for survival analyses. Rats who participated in the test (that is, had a latency to feed less than 15 min) were considered “death events” while those who did not participate “survived”. We then analyzed this by the Kaplan-Meier estimator and log-rank test to examine the effect of stress exposure between each genotype.

*Krtcap3* expression between WT and KO adolescents in multiple tissues was assessed by a t-test per tissue, and p-values for multiple comparisons were adjusted via the Holm-Sidak method. This same method was applied to comparing *Krtcap3* expression between WT control and stress animals from the main study. Expression analyses of the other genes, which included KO rats, were first analyzed by a two-way ANOVA, and then appropriately deconstructed to examine effects of stress per genotype and effects of genotype per stress condition.

## Results

### Verification of the Krtcap3 knock-out in multiple tissues

We verified the *Krtcap3*-KO in multiple tissues in female adolescent rats (**Supplementary Figure 1**). Compared to WT, KO rats had significantly lower *Krtcap3* expression in all tissues that were assessed: pituitary (T_3.13_ = 11.95, p_adj_ = 6.17e-3), adrenal (T_4.77_ = 11.03, p_adj_ = 9.95e-4), liver (T_3.02_ = 6.94, p_adj_ =0.024), hypothalamus (T_3.33_ = 4.84, p_adj_ = 0.026), ileum (T_3.05_ = 8.34, p_adj_ = 0.017), ovary (T_3.03_ = 3.55, p_adj_ = 0.038), and RetroFat (T_5.46_ = 4.35, p_adj_ = 0.024) tissues.

### Chronic stress exposure increased early fat mass and cage-wide food intake in both genotypes, but effects on growth, individual food intake, and final adiposity are seen only in KO rats

There were no differences in body weight at weaning (three weeks of age) between WT or KO rats, and no differences in weight between animals assigned to the control or the stress groups (**Supplementary Figure 2a**). Despite exposure to noise stress for three weeks, there were no differences in body weight by genotype nor by stress exposure at six weeks of age at HFD start (**Supplementary Figure 2b**). There was, however, a modest increase in total fat mass in noise-exposed rats compared to control rats for both genotypes (F_1, 26_ = 5.38, p = 0.03; **Supplementary Figure 2c**).

Contrary to what we had hypothesized, <u>the entirety of the stress protocol</u> described here increased the body weight of the rats over time (F_2.42, 67.89_ = 3.13, p = 0.041) rather than decreased it. When WT and KO growth curves are assessed separately, however, stress exposure ultimately does not impact WT body weight over time (**Figure 2**), while it does increase KO body weight over time (F_22, 308_ = 2.03, p = 0.005; **Figure 2**).

**Figure 2.**
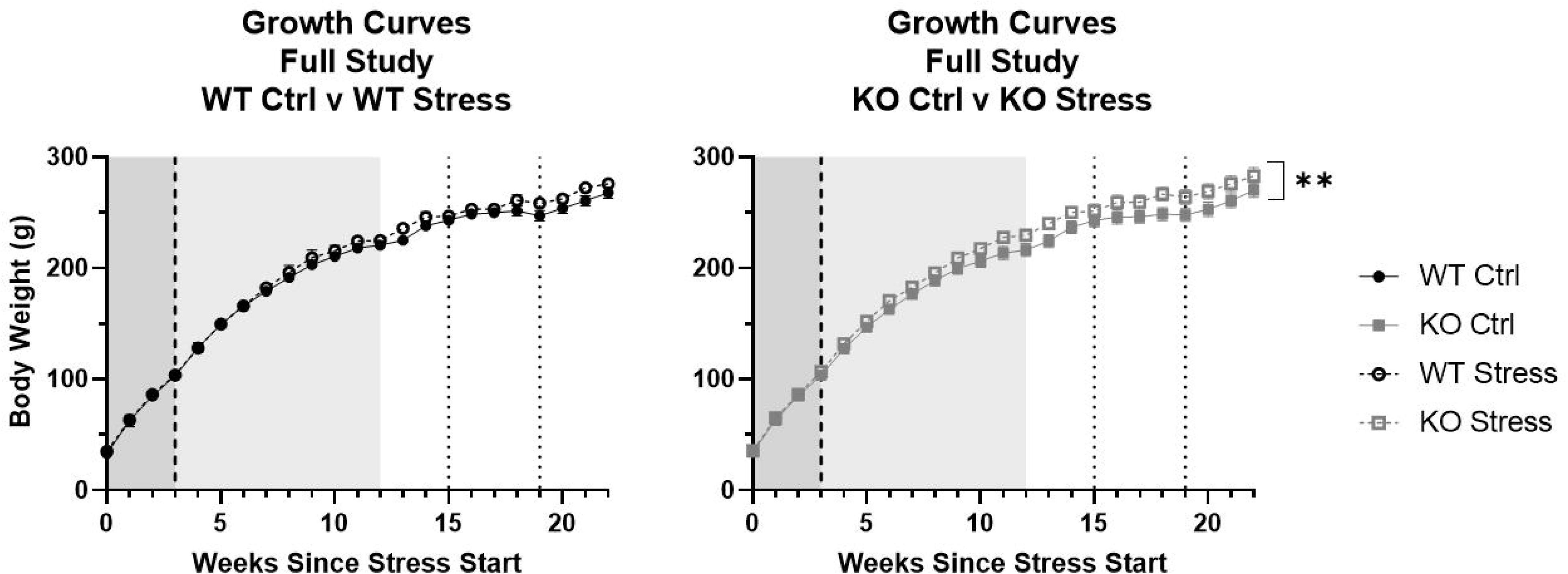
Stress exposure increased growth in *Krtcap3* knock-out (KO, gray square) rats but not in wild-type (WT, black circle) rats. Data are split by genotype to assess differences due to stress exposure respective for WT and KO rats. The mild stress period is demarcated by the shaded region from weeks 0 to 3 (white noise, dark gray) and from weeks 3 to 12 (white noise + additional mild stress, light gray), while the dashed lines at weeks 15 and 19 indicate the beginning and end of UCMS. High fat diet start is shown by the dashed line three weeks after the start of stress. The full growth curve of the 22-week long study demonstrates that control (filled) and stress-exposed (empty) WT rats had similar body weights for much of the study yet stress-exposed KO rats gain more weight over time compared to control KO rats. **p < 0.01 represents an interaction between body weight and time in the KO rats.

During the <u>mild stress protocol</u> (weeks 0-12), there was a slight increase in average weekly cage food consumption in the stress-exposed rats of both genotypes (F_1, 12_ = 4.41, p = 0.058; **Figure 3a**). EchoMRI analysis performed 13 weeks after stress initiation showed that stress-exposed rats of both genotypes had increased total fat mass relative to control counterparts (F_1, 27_ = 16.18, p = 4.17e-4; **Figure 3b**) as well as increased total lean mass (F_1, 27_ = 8.74, p = 6.4e-3; **Figure 3c**).

**Figure 3.**
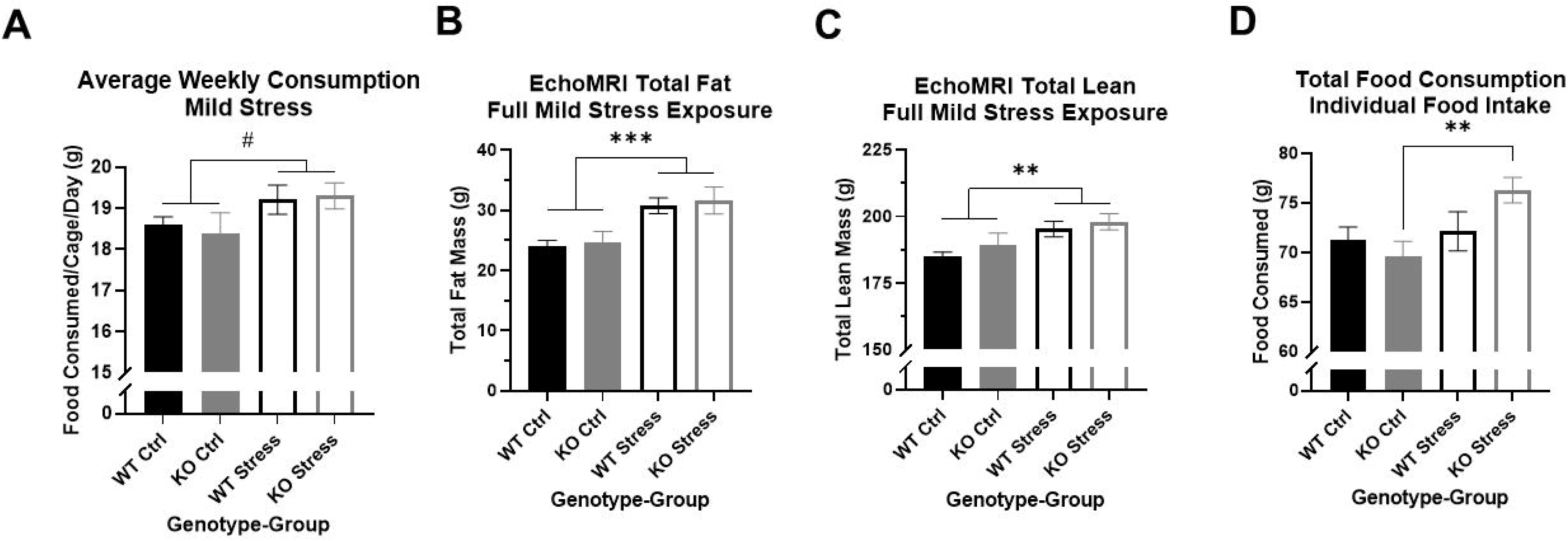
Mild stress exposure increased adiposity in wild-type (WT, black) and *Krtcap3* knock-out (KO, gray) rats without strongly impacting cage-wide eating behavior, but individual housing only increased eating in KO rats. (A) Stress-exposure mildly increased cage-wide food consumption in both WT and KO rats. #p < 0.1 represents main effect of stress exposure. (B) At the end of the mild stress protocol, EchoMRI analysis assessed total fat mass, and found that mild stress exposure had significantly increased total fat mass in both WT and KO rats. ***p < 0.001 represents main effect of stress. (C) Similarly, mild stress exposure significantly increased total lean mass in both WT and KO rats. **p < 0.01 represents main effect of stress. (D) Contrary to the preceding weeks, during the week of individual housing, stress-exposed KO rats consumed a greater quantity of food during the week, with no differences in WT eating behavior related to stress exposure. **p < 0.01 represents effect of stress in KO rats.

Within the <u>single week of individual housing</u> there was a main effect of stress exposure on the sum of food consumed per rat (F_1, 27_ = 5.53, p = 0.026) but also a nearly significant interaction between genotype and stress exposure (F_1, 27_ = 3.44, p = 0.075). Between WT rats there were no differences in the total amount of food consumed during the week, but stress-exposed KO rats consumed a larger quantity of food compared to control counterparts (T_13_ = 3.3, p = 5.53e-3; **Figure 3d**).

Four weeks of <u>UCMS exposure</u> did not alter the patterns we had seen during the earlier mild stress protocol. From start to end of UCMS (weeks 15-19), stress-exposed WT and KO rats gained more weight than control counterparts (F_1, 28_ = 435.1, p = 1.61e-5; **Supplementary Figure 3a**). Despite this difference, UCMS exposure only mildly increased average weekly cage food consumption (F_1, 12_ = 3.68, p = 0.075; **Supplementary Figure 3b**) in both genotypes. EchoMRI analysis conducted at the conclusion of UCMS exposure determined that stress exposure increased total fat mass (F_1, 28_ = 5.58, p = 0.025), though this was driven by differences in the KO rats (T_14_ = 2.34, p = 0.035; **Supplementary Figure 3c**). Stress exposure also increased total lean mass (F_1, 27_ = 4.34, p = 0.047; **Supplementary Figure 3d**).

At the <u>end of the study</u>, stress-exposed rats were only slightly heavier than control rats, for both genotypes (F_1, 28_ = 3.46, p = 0.074; **Figure 4a**). Similar to the post-UCMS findings, there was a main effect of stress for RetroFat mass (F_1, 26_ = 5.43, p = 0.028), that was driven by the KO rats (T_12_ = 2.58, p = 0.024; **Figure 4b**) with no differences in WT rats. Stress had a modest impact on ParaFat (F_1, 27_ = 3.52, p = 0.071; **Figure 4c**) and OmenFat (F_1, 28_ = 3.52, p = 0.071; data not shown). There were no differences in body length, tail length, or organ weight by genotype nor by stress exposure (data not shown). Control KO rats had smaller adrenal glands compared to control WT rats (T_13_ = 2.07, p = 0.059; **Figure 4d**), although there were no differences in adrenal gland weight between stress-exposed WT and KO rats. While there is a visual increase in adrenal gland weight between control and stress-exposed KO rats, due to large variation this was not statistically significant.

**Figure 4.**
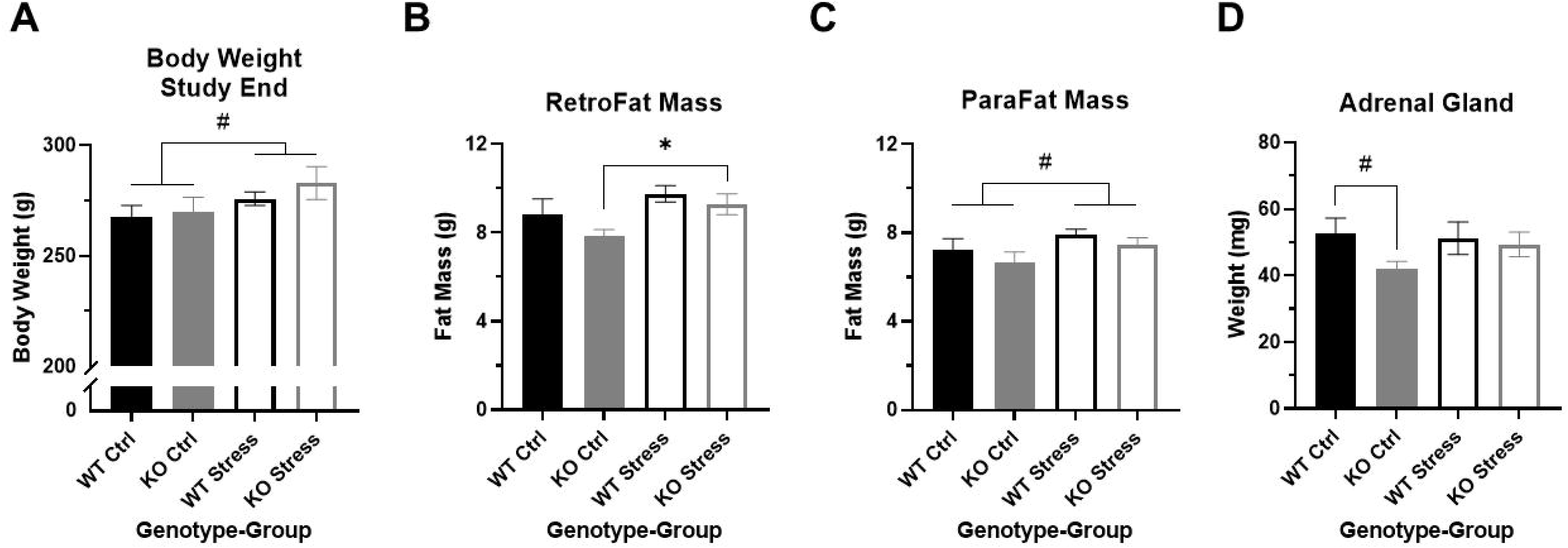
Stress exposure increased retroperitoneal fat (RetroFat) mass in *Krtcap3* knock-out (KO, gray) rats but not wild-type (WT, black) rats. (A) Stress-exposed (empty) rats of both genotypes had a slightly greater body weight at study conclusion compared to control (filled) counterparts. #p < 0.1 represents main effect of stress. (B) Stress-exposed KO rats had increased RetroFat mass compared to control counterparts, with no differences between WT rats. *p < 0.05 represents effect of stress for KO rats. (C) There was a mild main effect of stress exposure on parametrial fat (ParaFat) mass, where stress-exposed rats of both genotypes had increased fat pad mass relative to control counterparts. #p < 0.1 represents main effect of stress. (D) KO rats naïve to stress had slightly smaller adrenal glands compared to control WT rats. #p < 0.1 represents effect of genotype for control rats.

### No differences in fasting glucose or glucose tolerance by genotype nor by stress exposure

There were no differences in fasting glucose or glucose response to a glucose challenge, neither by genotype nor by stress exposure (data not shown).

### Initial mild stress increased NSF anxiety-like behaviors in KO, but not WT

There was a significant interaction between genotype and stress exposure regarding the overall movement of the rat during the NSF, as measured by the number of line crossings (F_1, 28_ = 13.22, p = 1.1e-3). When we assessed each genotype separately, we found that there was little difference in activity between WT groups (T_14_ = 1.78, p = 0.096; **Figure 5a**), but stress-exposed KO rats moved much less than their control counterparts (T_14_ = 3.35, 4.8e-3; **Figure 5a**). There were similar results in the number of center approaches the rats made, with a significant interaction between genotype and stress exposure (F_1, 26_ = 6.23, p = 0.019) where stress-exposed KO rats approached much less frequently than control KO rats (T_13_ = 3.78, p = 2.3e-3; **Figure 5b**), with no significant differences between WT rats. When measuring the amount of time spent in the center there was a main effect of stress group, where control rats of both genotypes were more willing to spend time in the center of the box compared to stress-exposed rats (F_1, 26_ = 7.88, p = 9.4e-3; **Figure 5c**).

**Figure 5.**
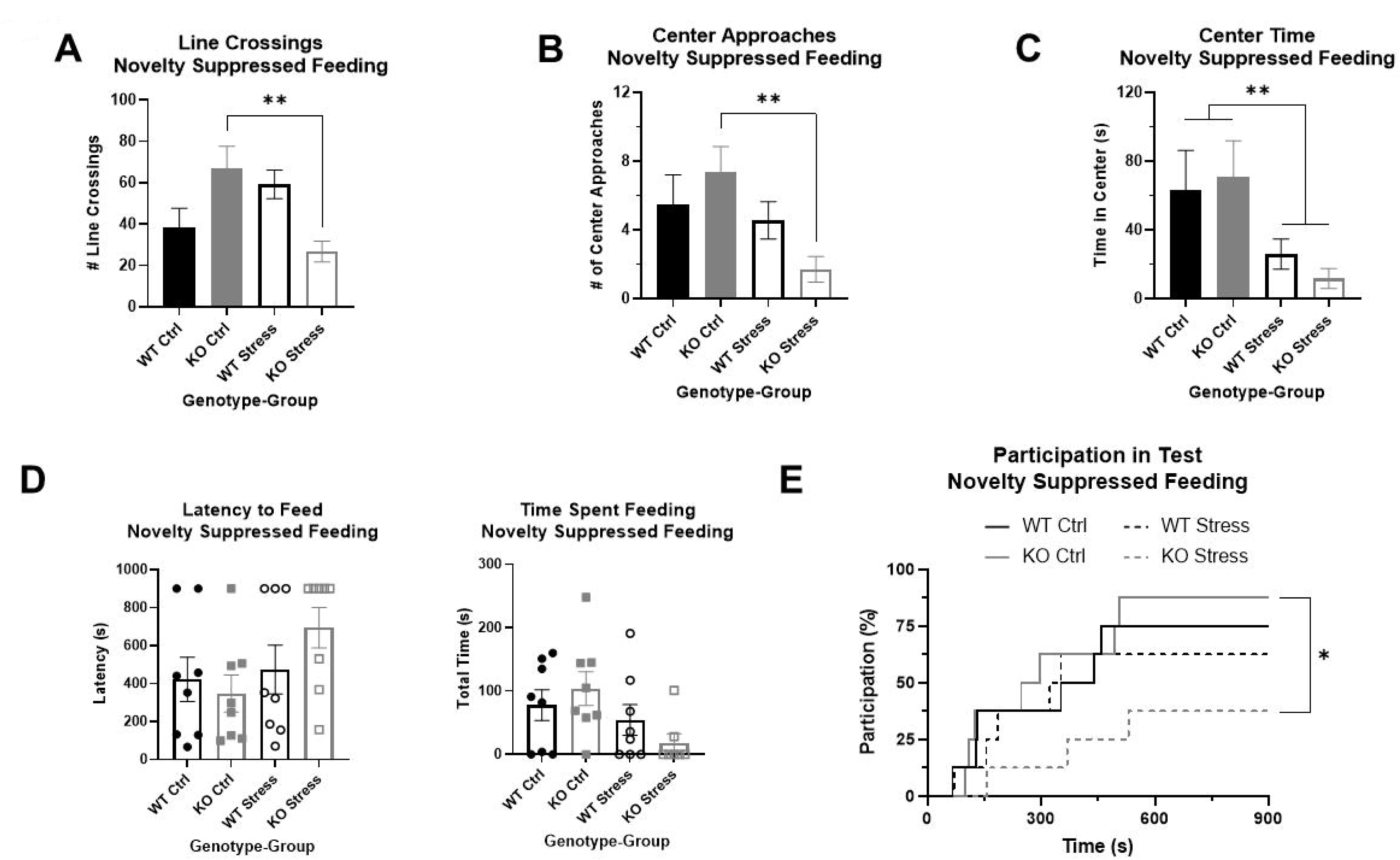
Novelty suppressed feeding test demonstrates that stress exposure did not greatly alter anxiety in wild-type (WT, black) rats, but did significantly increase anxiety in *Krtcap3* knock-out (KO, gray) rats. (A) There was a significant interaction between genotype and group for number of line crossings where stress-exposed (empty) KO rats moved much less than control counterparts (filled) with no changes in WT rats. **p < 0.01 represents effects of stress respective to each genotype. (B) There were no differences in frequency of center approaches between control and stress WT rats, but stress-exposed KO rats approached the center fewer times than controls. **p < 0.01 represents effect of stress for KO rats. (C) For both WT and KO rats, stress-exposed rats spent much less time in the center of the field compared to controls. **p < 0.01 represents a main effect of stress. (D) When directly evaluating latency to feed and time spent feeding, there were ceiling and floor effects that precluded statistical analysis by a two-way ANOVA. Visually, there is little difference between control and stress-exposed WT rats (circle), but a large difference in KO rats (squares). Instead, we re-conceptualized these phenotypes as test participation instead. (E) Stress exposure (dashed line) had little effect on the probability of participation of WT rats compared to control counterparts (solid line), but KO rats exposed to stress were much less likely to consume food during the test compared to controls. *p < 0.05 represents effect of stress for KO rats.

Because of ceiling and floor effects, we could not accurately measure statistical differences in latency to feed or the total time a rat spent feeding (**Figure 5d**). We instead compiled these phenotypes into one that examined participation in the test and used modified survival curves to analyze the likelihood of a rat participating given its genotype and stress exposure. There were no significant differences in WT participation in the test due to stress exposure, but stress-exposed KO rats had a significantly lower likelihood of participating in the NSF than controls (Χ^2^ = 5.3, p = 0.02; **Figure 5e**).

### Initial mild stress increased FST stress response only in KO rats

We evaluated immobility and mobility in the FST on Day 2, plus the change in immobility from the first five minutes of Day 1 to Day 2, as measures of passive coping response to stress. There were main effects by stress exposure for both immobility and mobility on Day 2: as expected, stress exposure increased immobility in the test (F_1, 27_ = 10.21, p = 3.5e-3; **Figure 6a**) and decreased mobility (F_1, 27_ = 10, p = 3.8e-3; **Figure 6b**). In addition, there was a strong effect of stress exposure on the change in immobility between the days (F_1, 26_ = 24.04, p = 4.34e-5; **Figure 6c**), where stress-exposed rats had a smaller change between the days. Importantly, there was a main effect of genotype (F_1, 26_ = 4.25, p = 0.049) driven by differences in the stress-exposed rats (T_13_ = 3.19, p = 7.2e-3; **Figure 6c**): KO rats had significantly greater immobility on Day 2 relative to Day 1, while WT rats had a slight decrease in immobility on Day 2 compared to Day 1. These differences support an increased response to FST stress in the KO rats exposed to mild stress relative to WT rats.

**Figure 6.**
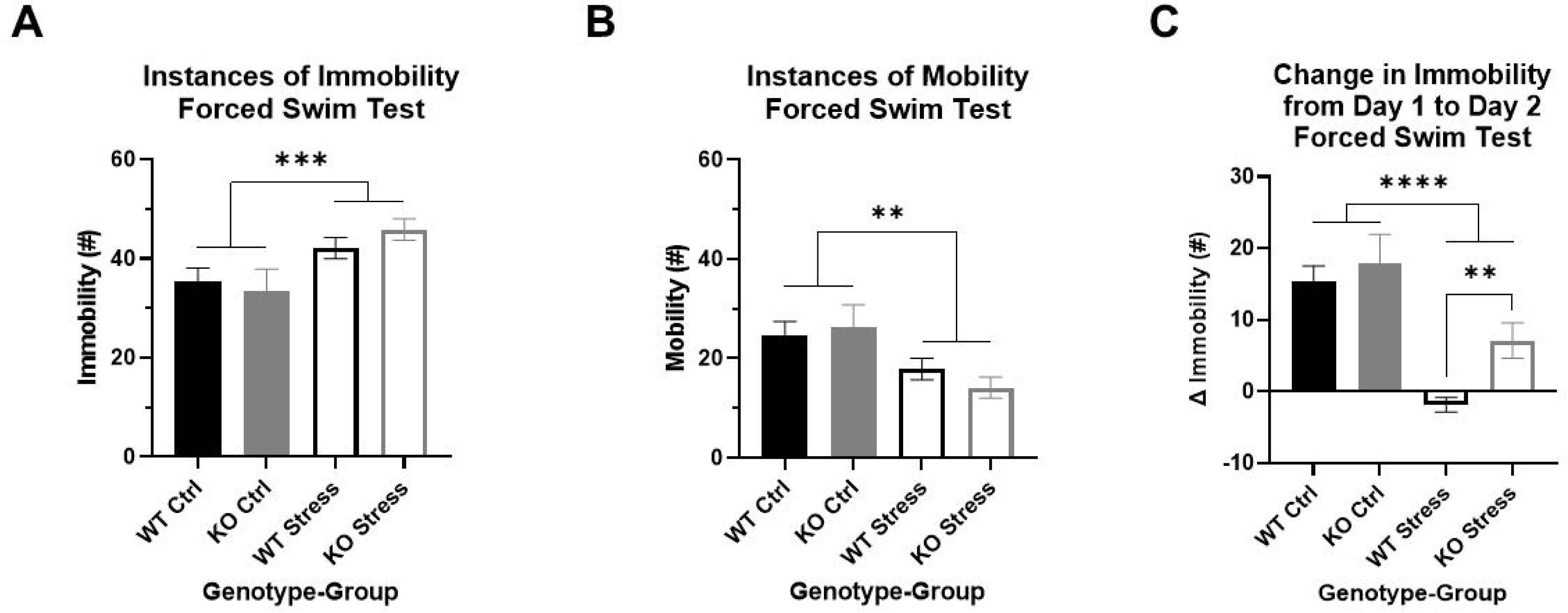
Forced swim test indicates that stress exposure more greatly alters stress coping in *Krtcap3* knock-out (KO, gray) rats compared to wild-type (WT, black) rats. The FST is a two-day test: Day 1 is a 15-min training swim while Day 2 is a 5-min test swim. (A) When compared to control rats (filled), stress-exposed rats (empty) of both genotypes had increased immobility and (B) decreased mobility compared to control rats (filled) on Day 2 of the test. **p < 0.01 represents a main effect of stress. (C) The change in immobility from Day 1 to Day 2 was significantly greater in control rats compared to stress-exposed rats, regardless of genotype. However, within the stress-exposed rats, immobility did not change in WT rats but had continued to increase between the days in KO rats, indicating differences in the stress coping response. **p < 0.01 represents effect of genotype respective to stress exposure; ****p < 0.0001 represents a main effect of stress.

### KO rats are more exploratory in OFT, and show decreased center entries in response to stress, relative to WT

After UCMS exposure, we used an OFT to measure locomotor and anxiety-like behavior in the rats. Over the full course of the test, KO rats spent more time moving compared to WT counterparts, regardless of stress exposure (F_1, 27_ = 4.23, p = 0.049; **Figure 7a**). Additionally, stress-exposed rats of both genotypes spent more time moving than controls (F_1, 27_ = 4.23, p = 0.049; **Figure 7a**). KO rats reared more relative to WT rats (F_1, 28_ = 4.49, p = 0.043), though this was primarily in the control rats (T_14_ = 2.65, p = 0.019; **Figure 7b**) rather than the stress-exposed rats. KO rats also tended to travel a greater distance (F_1, 27_ = 4.09, p = 0.053; **Figure 7c**) than WT counterparts under both control and stress conditions.

**Figure 7.**
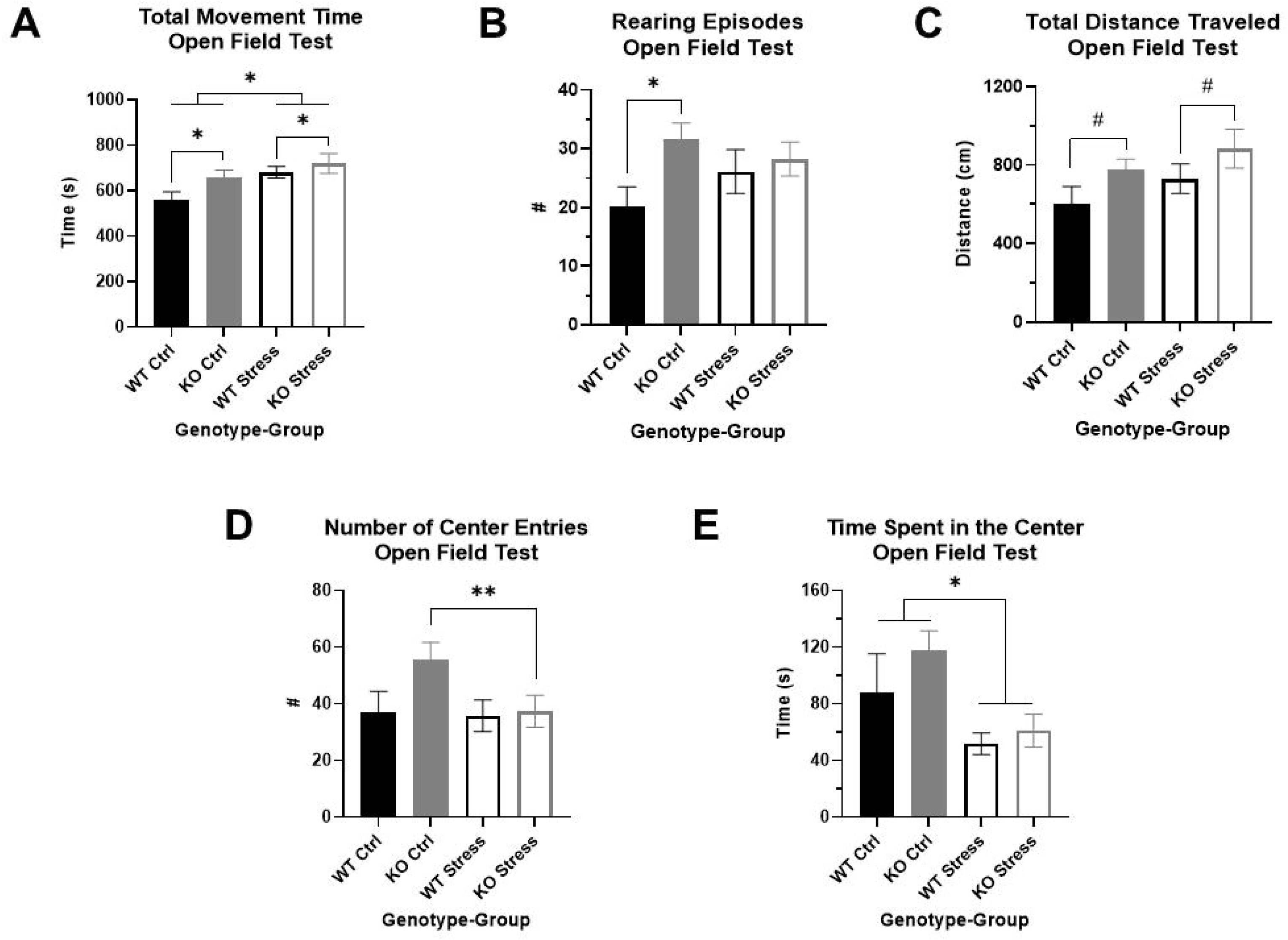
Open field test indicates that *Krtcap3* knock-out (KO, gray) rats are more exploratory than wild-type (WT, black) rats. (A) KO rats spent more time moving compared to WT counterparts, plus stress-exposed (empty) rats of both genotypes spent more time moving compared to respective control counterparts (filled). *p < 0.05 represents a main effect of genotype and a main effect of stress. (B) Overall, KO rats of both stress conditions reared more frequently than WT rats, demonstrating increased vertical exploration. *p < 0.05 represents a main effect of genotype. (C) Additionally, KO rats of both stress conditions travelled a slightly greater distance in the field compared to WT rats, indicating increased horizontal exploration. #p < 0.1 represents a main effect of genotype. (D) While the two-way ANOVA did not pick up an interaction between genotype and stress exposure, an examination of the genotypes separately revealed that stress exposure did not alter WT willingness to approach the center of the box, but stress-exposed KO rats approached much less frequently than control counterparts. **p < 0.01 represents an effect of stress for KO rats. This is consistent with results from the novelty suppressed feeding test, signifying that stress exposure more strongly impacted anxiety-related behaviors in KO rats than WT rats. (E) Also consistent with prior behavioral results, stress exposure for both genotypes did decrease amount of time spent in the center of the box. *p < 0.05 represents main effect of stress.

While the two-way ANOVA did not show an interaction between genotype and stress exposure that impacted number of center entries, we previously saw in the NSF that stress affected willingness to approach the center in KO rats but not WT. When we examined each genotype separately, we found that there were no differences by stress exposure in WT rats, but stress-exposed KO rats entered the center of the box much less frequently than control counterparts (T_13_ = 3.78, p = 2.3e-3; **Figure 7d**). As we had previously seen in the NSF, stress exposure affected amount of time spent in the center, where stress-exposed rats of both genotypes spent less time in the center than controls (F_1, 27_ = 4.41, p = 0.045; **Figure 7e**). Neither genotype nor stress exposure affected distance traveled in the center of the box, however (data not shown).

### Mild stress exposure did not increase CORT in stress-exposed rats relative to control counterparts in both WT and KO rats

After 10 weeks of mild stress exposure, rats were administered an acute restraint test. There were no statistically significant differences in basal CORT by genotype nor by stress exposure (**Figure 8a**), although KO rats visually have a slightly lower basal CORT than WT. As restraint continued, CORT increased in all rats regardless of genotype or stress exposure (F_2, 40_ = 317.23, p = 2.9e-25), both after 10 (Q_25_ = 20.07, p < 1e-4; **Figure 8b**) and 30 (Q_27_ = 20.61, p < 1e-4; **Figure 8b**) minutes of restraint. When we examined the effect of genotype and stress exposure separately for each time point, there was a significant interaction after 10 minutes of restraint (F_1, 26_ = 4.73, p = 0.039) where stress-exposed KO rats had a lower CORT than control counterparts (T_13_ = 2.29, p = 0.039; **Figure 8c**), with no differences in WT rats. Although stress-exposed KO rats maintain a visually lower CORT after 30 minutes of restraint compared to control KO rats, there was no longer a statistically significant difference.

**Figure 8.**
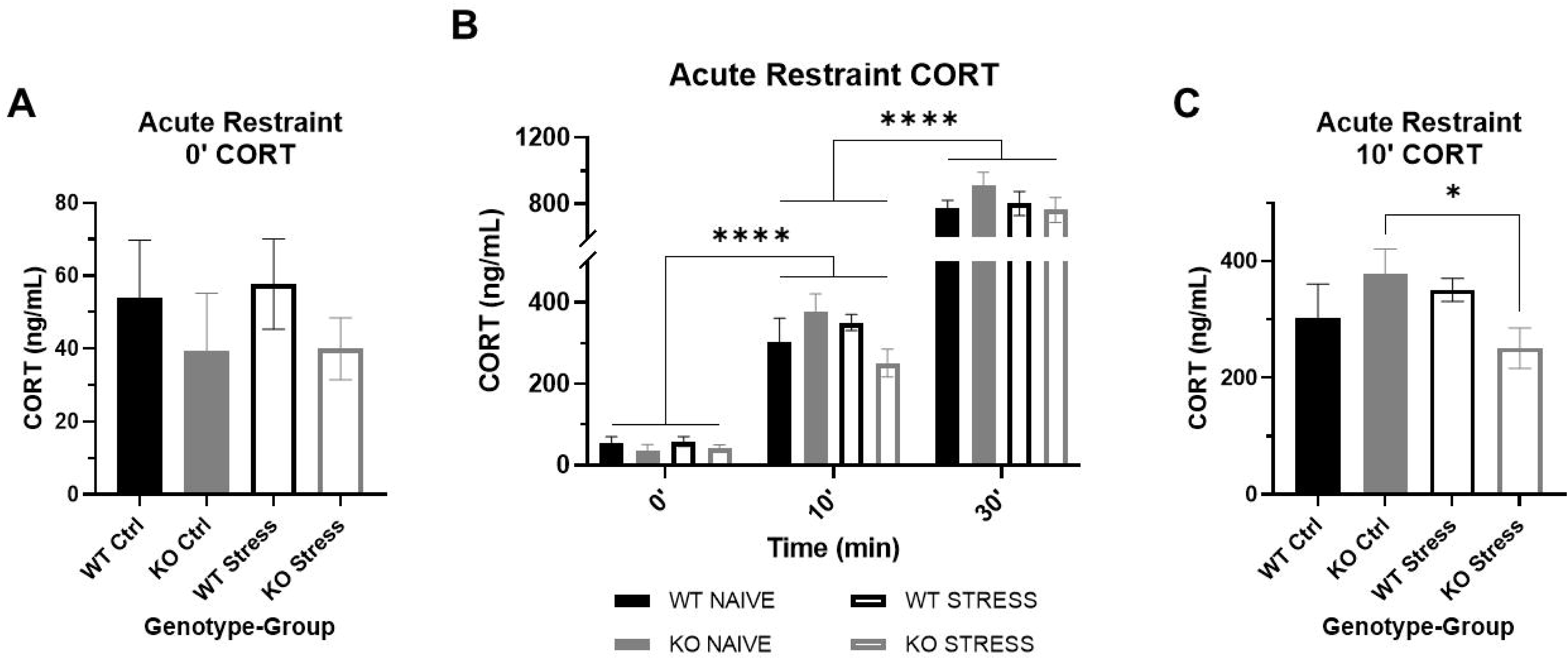
Corticosterone (CORT) response to acute restraint stress in wild-type (WT, black) and *Krtcap3* knock-out (KO, gray) rats. (A) Serum basal CORT was measured immediately prior to acute restraint after 10 weeks of the mild stress protocol. Basal CORT was not increased in stress-exposed rats (empty) relative to control counterparts (filled). (B) Serum CORT measurements were also taken after 10 and 30 minutes of restraint. CORT significantly increased as restraint continued in all rats. ****p < 0.0001 represents effect of time for all groups. (C) After 10 minutes of restraint, there were no differences in CORT between WT rats, but stress-exposed KO rats had a lower CORT than control KO rats. *p < 0.05 represents effect of stress exposure in KO rats.

Prior to UCMS, we measured serum CORT in the rats and saw no differences by genotype nor by stress exposure (data not shown). This confirmed that the mild stress protocol did not increase basal CORT in WT rats and indicates that the changes in adiposity and behavior in KO rats occurred despite lack of changes in basal CORT.

### At the end of the study, CORT is lower in KO vs WT under control conditions, and increases in response to stress in KO but not WT rats

We also measured CORT collected at euthanasia. Given prior results (Szalanczy, Giorgio et al. 2023), we expected that there would be an effect of euthanasia order on CORT, so we initially examined only the first rat of each cage, presumed to be the basal measurement. There was a significant interaction between genotype and stress exposure (F_1, 12_ = 9.93, p = 0.008), where stress-exposed KO rats had elevated CORT relative to control counterparts (T_4.67_ = 2.91, p = 0.036; **Figure 9a**) yet contrary to expectation stress-exposed WT rats had slightly lower CORT relative to control counterparts (T_5.98_ = 2.11, p = 0.08). We do not fully understand why basal CORT was so high in control WT rats and these results may be confounded by an unknown stress at the time of sac. Similar to our previous work (Szalanczy, Giorgio et al. 2023), we also found that control KO rats have significantly lower serum CORT relative to control WT rats at the time of euthanasia (T_3.59_ = 3.31, p = 0.035; **Figure 9a**), indicating KO rats have lower basal stress compared to WT rats, supporting the behavioral studies described above.

**Figure 9.**
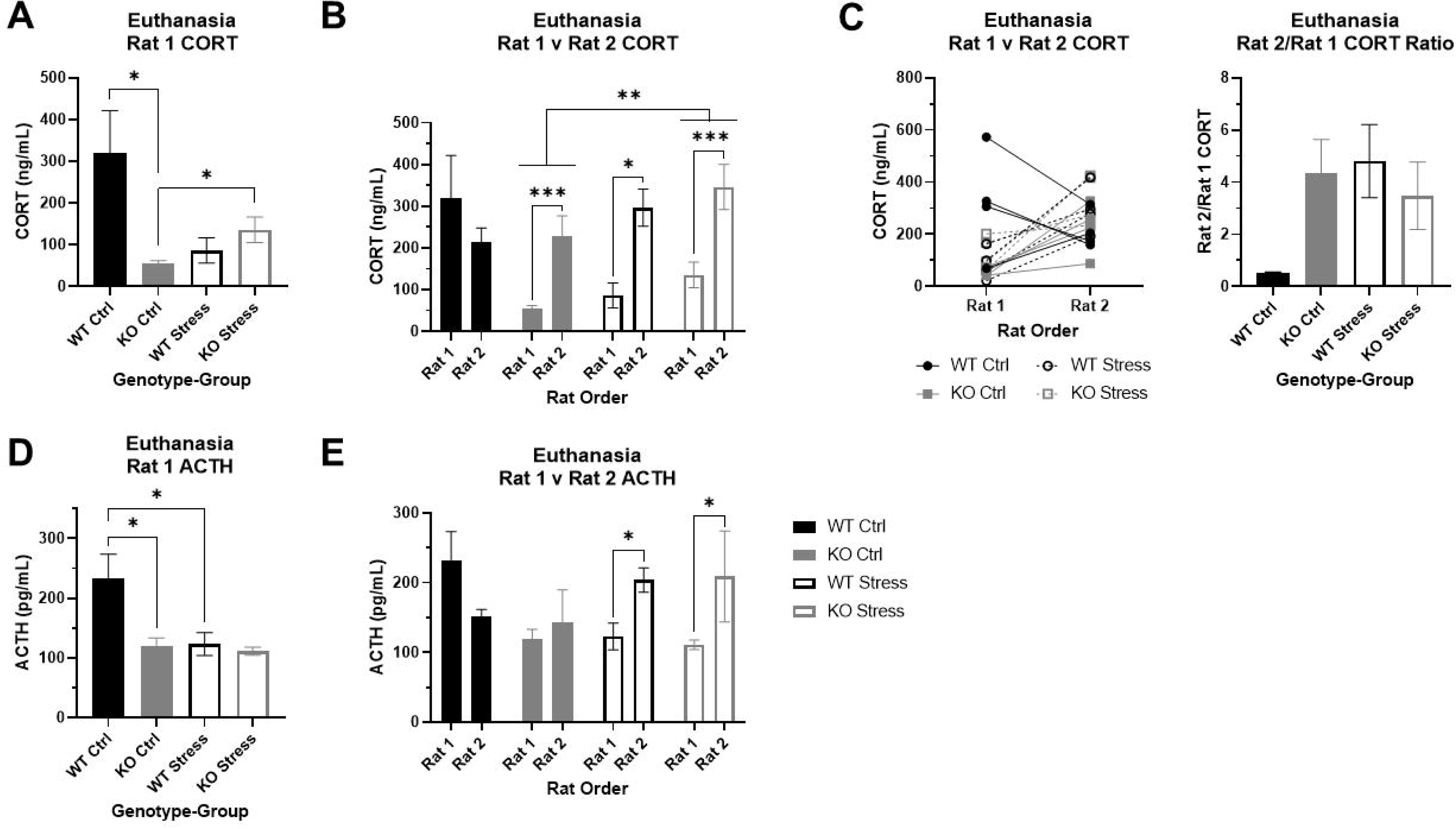
Corticosterone (CORT) and adrenocorticotropic hormone (ACTH) at euthanasia in wild-type (WT, black) and *Krtcap3* knock-out (KO, gray) rats. Serum was collected from trunk blood at euthanasia, and rats were euthanized within the cage one at a time, with Rat 1 euthanized first and Rat 2 euthanized second. (A) Control (filled) KO rats had lower basal CORT compared to control WT rats. Additionally, CORT of KO Rat 1 was greater in those that were stress-exposed (empty) compared to control rats, but not for WT rats. *p < 0.05 represents effect of genotype in control rats and effect of stress exposure in KO rats. (B) We confirmed that stress exposure increased CORT in KO rats, for both Rat 1 and Rat 2. **p < 0.01 represents effect of stress exposure in KO rats. Additionally, we found that KO rats show a significant increase in CORT when their cage-mate is removed at euthanasia regardless of stress exposure; we found the same pattern in stress-exposed WT rats, but not control WT rats. *p < 0.05 represents effect of euthanasia order in WT rats, ***p < 0.001 represents effect of euthanasia order in KO rats. (C) The first figure demonstrates the difference in CORT between Rat 1 and Rat 2 of each cage, where WT rats are represented by black circles and KO rats by gray squares. Control rats are marked with filled symbols and a solid line, while stress-exposed rats have empty symbols and dashed lines. Examination of the CORT ratio between Rat 1 and Rat 2 for each genotype and stress condition confirmed that stress exposure does not alter this ratio in KO rats, but does increase the ratio in WT rats. #p < 0.1 represents effect of stress for WT rats. (D) ACTH was lower in control KO Rat 1 compared to control WT rats. Further, ACTH was lower in stress-exposed WT Rat 1 relative to control counterparts. *p < 0.05 represents effect of genotype in control rats and effect of stress exposure in WT rats. (E) We found that ACTH is significantly increased in Rat 2 for stress-exposed rats, whether WT or KO. There were no differences by order in control rats. *p < 0.05 represents effect of order in stress-exposed rats.

### Euthanasia order alters serum CORT in both control and stress-exposed KO rats as well as stress-exposed WT rats

We next assessed the impact of euthanasia order in addition to genotype and stress exposure. Similar to our previous work (Szalanczy, Giorgio et al. 2023), there was a significant effect of order (F_1, 24_ = 21.01, p = 1.2e-4), with rats euthanized second showing higher serum CORT. There was still an interaction between genotype and stress exposure (F_1, 24_ = 8.65, p = 7e-3), but there was also a three-way interaction between genotype, stress exposure, and euthanasia order (F_1, 24_ = 6.6, p = 0.017). We split the data by genotype to confirm the impact of stress exposure and order on CORT, and found a significant main effects of stress (F_1, 12_ = 9.53, p = 9e-3; **Figure 9b**) and order (F_1, 12_ = 29.72, p = 1.47e-4; **Figure 9b**) in KO rats, demonstrating that chronic stress exposure increased CORT in the KO rats and that CORT rises in the second rat of the cage regardless of prior stress exposure. In WT rats, we found an interaction between stress exposure and order (F_1, 12_ = 6.28, p = 0.028) where order did not impact CORT in control WT rats but CORT was significantly elevated in the second rat of the cage in stress-exposed WT rats (T_5.25_ = 3.91, p = 0.01; **Figure 9b**).

These findings were supported by additional analyses of the CORT ratio between the first and second rat of the cage for each genotype and stress condition. There was a nearly significant interaction between genotype and stress exposure (F_1, 11_ = 4.21, p = 0.065) where WT control rats had a much lower ratio relative to the other three groups (**Figure 9b**). These data indicate a stronger social stress response to removal of cage-mate in KO rats, which is seen only in WT rats when they have previously been exposed to chronic stress.

### Basal ACTH is lower in KO vs WT rats under control conditions

Given that there was a significant effect of euthanasia order on CORT, we initially examined ACTH in only the first rats of the cage. There were main effects of genotype (F_1, 12_ = 5.83, p = 0.033) and stress exposure (F_1, 12_ = 5.94, p = 0.031) where control WT rats had greater basal ACTH than control KO rats (T_4.98_ = 3.02, p = 0.03; **Figure 9d**) and stress-exposed WT rats (T_5.99_ = 2.46, p = 0.049; **Figure 9d**).

### ACTH increases in response to separation at euthanasia only in stress-exposed rats, regardless of genotype

When we incorporated data from the second rats and the factor of euthanasia order into the analyses, there was an interaction between stress exposure and order (F_1, 24_ = 5.76, p = 0.025) separate from genotype. Specifically, in control rats there were no differences in ACTH between the two rats of the cage, whereas in stress-exposed rats the second rat of the cage had elevated ACTH compared to the first rat (T_1, 14_ = 8.71, p = 0.011; **Figure 9e**).

### Expression of the GR Nr3c1 changes between genotypes and stress exposure in pituitary, liver, and colon

We investigated *Nr3c1* expression in the pituitary, colon, and liver. The pituitary and colon both have high *Krtcap3* expression (Szalanczy, Giorgio et al. 2023) and each are involved in stress response (Herman, McKlveen et al. 2016, Wiley, Higgins et al. 2016), indicating that they may be the tissues of action for *Krtcap3*. Further, we also sought to verify previously reported changes in *Nr3c1* expression in the liver in response to stress (Szalanczy, Giorgio et al. 2023). We found a very nearly significant main effect of stress-exposure on *Nr3c1* expression in the pituitary (F_1, 27_ = 4.15, p = 0.052), where stress-exposed KO rats had greater *Nr3c1* expression in the pituitary than control counterparts (T_14_ = 2.33, p = 0.035; **Figure 10a**) with no differences in expression between WT rats. Similarly, there was a near interaction in colon *Nr3c1* expression (F_1, 25_ = 4.2, p = 0.051) where there was no difference by stress exposure in WT rats, but stress exposure slightly decreased *Nr3c1* expression in KO rats (T_13_ = 2.06, p = 0.06; **Figure 10b**). In the control condition, KO rats also had higher colon *Nr3c1* expression than WT rats (T_13_ = 2.2, p = 0.047; **Figure 10b**). These findings mimic the metabolic and behavioral findings, where differences are seen between control and stress-exposed KO rats but not between stress conditions in WT rats, further supporting an important role for *Krtcap3* in the stress axis. As expected, *Nr3c1* expression was also increased in the liver tissue of stress-exposed rats (F_1, 27_ = 4.51, p = 0.043), but the difference here was driven by the WT rats (T_13_ = 3.11, p = 8.2e-3; **Figure 10c**) with no changes in the KO rats. Similar to our findings in the colon, we also identified a difference in liver *Nr3c1* expression between control WT and KO rats, where KO rats had higher expression (T_13_ = 2.26, p = 0.041; **Figure 10c**). On the other hand, there were no differences in pituitary *Pomc* expression or colon *Hsd11*β*2* by genotype or by stress exposure (data not shown).

**Figure 10.**
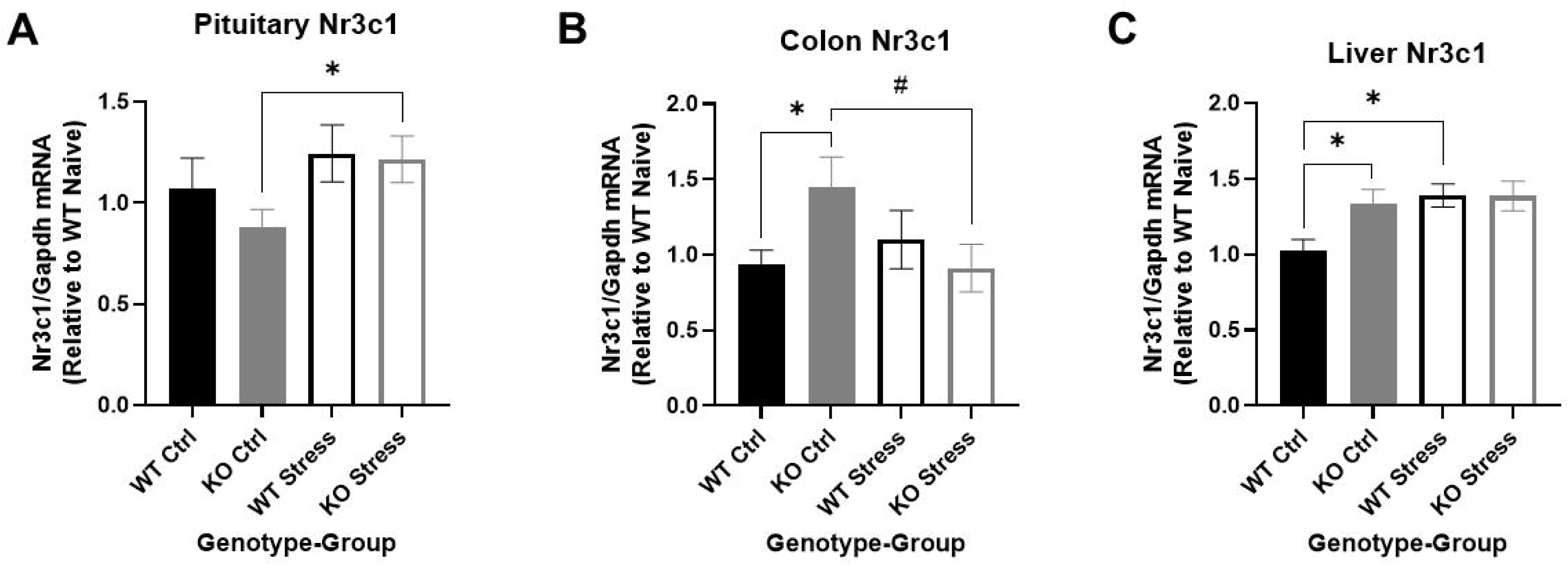
Differential expression of the glucocorticoid receptor *Nr3c1* by genotype and stress exposure in pituitary, colon, and liver. We compared *Nr3c1* expression between wild-type (WT, black) and *Krtcap3* knock-out (KO, gray) control rats (filled) and exposed to stress (empty) in multiple tissues. (A) Stress-exposed KO rats had greater pituitary *Nr3c1* expression than control counterparts, with no significant difference in WT rats. *p < 0.05 represents effect of stress for KO rats. (B) In the colon, control KO rats had greater *Nr3c1* expression than control WT and greater expression than stress-exposed KO rats. #p < 0.1 represents effect of stress for KO rats, *p < 0.05 represents effect of genotype for control rats. (C) In the liver, on the other hand, control WT rats had lower *Nr3c1* expression than control KO rats and stress-exposed WT rats. Stress exposure did not alter expression in KO rats. *p < 0.05 represents effect of stress for WT rats and effect of genotype for control rats.

### Krtcap3 mRNA expression does not change between control and stress-exposed WT rats

We measured *Krtcap3* expression between control and stress-exposed WT rats in key tissues to evaluate if stress exposure altered expression. Ultimately, there were no differences in *Krtcap3* expression between control and stress-exposed WT rats in the pituitary, the adrenal, or the colon (data not shown).

## Discussion

We demonstrate here that chronic stress impacted both metabolic and behavioral phenotypes to a greater extent in KO rats than in WT rats. Although the differences were mild, stress-exposed KO rats gained more weight, ate more food when isolated, and had greater fat mass relative to control counterparts. Stress-exposed KO rats also exhibited increased anxiety-like behavior and had a greater passive coping response than control KO rats. In contrast, stress-exposed WT rats showed minimal differences in all of these measures compared to control WT rats. These data support an important role for *Krtcap3* in the stress response and endorse that its impact on adiposity is influenced by stress. Similar to previous work (Szalanczy, Giorgio et al. 2023), KO rats in either stress condition demonstrate a strong CORT response when their cage-mate is removed at euthanasia, demonstrating that KO rats are more sensitive to psychosocial stress. In addition, we found that chronic stress altered *Nr3c1* expression in pituitary and colon only in the KO rats, while control KO rats have higher *Nr3c1* expression in the colon and liver compared to WT. While more work remains to be done to elucidate the role of *Krtcap3* in the stress pathway, this work is the first to demonstrate that *Krtcap3* plays a role in the stress response of WKY rats and that this has downstream impacts on adiposity and behavior.

Current findings indicate that the stress protocol in this study differs from conditions we have previously reported (Szalanczy, Goff et al. 2022, Szalanczy, Giorgio et al. 2023) and this may be because the noise we utilized here was not powerful enough to recapitulate construction vibrations (Reynolds, Li et al. 2018, Terashvili, Kozak et al. 2020) that were likely present in the first study. It is also possible that the lack of metabolic, behavioral, and CORT differences between WT control and stress-exposed rats are because 1) early mild stress led to desensitization of future stressors only in WT rats (Xu, Zhang et al. 2011, Mileva, Rooke et al. 2017, Morera, Gallea et al. 2020, Tran and Gellner 2023) or 2) WT rats could not distinguish between the chronic stress procedure and the stress of the study design (which included multiple behavioral and metabolic tests plus repeated blood collections). As WKY rats are known to be stress-sensitive (Redei, Udell et al. 2022), the latter option is more likely. Upcoming work from the Redei lab also suggests that WKY rats are chronically stressed when administered multiple behavioral tests despite long periods of rest in between (personal communication). If WT rats were not able to distinguish between the stress protocol and the study protocol, this work may indicate that low *Krtcap3* expression levels “normalize” HPA axis function and stress response in WKY rats. This is supported by 1) low basal stress, 2) the expected behavioral, metabolic, and CORT responses to chronic stress protocol, and 3) the expected changes in pituitary and colon *Nr3c1* expression levels in KO, but not WT, rats. Future studies that compare the *Krtcap3-*KO rats to other strains are needed to test this compelling hypothesis, and this work emphasized the importance of background strain and environmental conditions in rodent metabolic and behavioral studies.

### Stress induced greater changes in adiposity and eating behavior in KO rats than WT

As opposed to the suspected effect of construction noise in our previous study (Szalanczy, Goff et al. 2022), the chronic stress protocol here increased body weight over time only in the KO rats, with no effect in WT rats. Although stress exposure increased adiposity in both WT and KO rats early in the study, by the end the difference was only maintained in KO rats. In contrast to our previous work (Szalanczy, Giorgio et al. 2023), high basal CORT at euthanasia was associated with increased adiposity. This effect on adiposity more closely aligns with changes seen following psychosocial stress rather than physical stress (Patterson and Abizaid 2013) and may result from the number of times cage-mates were separated during the study. In support of this, when all rats were socially isolated, the stress-exposed KO rats ate significantly more food compared to control KO rats. We posit that prior stress-exposure increased susceptibility to the psychosocial stress of social isolation in KO rats, who may have increased food intake to ameliorate the anxiety-inducing effects of the separation (Maniam and Morris 2010, Kistenmacher, Goetsch et al. 2018, Hyldelund, Dalgaard et al. 2022), while WT rats were unaffected. Overall we show that KO rats responded differently to stress than WT rats, and that these differences influenced adiposity. Fully elucidating the difference in response between WT and KO rats to different stressors and understanding the impact on adiposity is a key goal of future studies.

### Stress exposure increased anxiety-like behaviors and passive coping only in KO rats

As we saw larger adiposity differences between control and stress-exposed KO rats, so too did we see larger behavioral changes in KO rats compared to WT, with greater increases in anxiety following stress exposure. Importantly, control KO rats exhibit lower anxiety measures in the OFT compared to WT rats, supportive of a lower basal stress state. In response to chronic stress, KO rats show the expected increases in thigmotaxis in the OFT as well as hyponeophagia and significant decreases in exploratory activity in the NSF. WT rats, on the other hand, displayed only minimal differences in these measures after stress exposure, in contrast to findings after chronic restraint stress in WKYs (Jung, Meckes et al. 2020). We also found that immobility in the FST increased in stress-exposed KO rats from Day 1 to Day 2 of the test, with minimal changes between days in stress-exposed WT rats. This suggests a greater passive coping response (Commons, Cholanians et al. 2017, Molendijk and de Kloet 2019) or greater depressive-like behavior (Redei, Udell et al. 2022) in KO rats following early life stress. As the WKY rat is hyper-sensitive to stress (Redei, Udell et al. 2022), these findings support that WT rats were unable to distinguish between the stress of the study design (control rats) and the chronic stress procedures (stress-exposed rats): both groups of rats were chronically stressed. The fact that KO rats show the expected changes in emotional behaviors in response to chronic stress suggests that low *Krtcap3* expression may normalize the stress response in WKY rats, although more work is needed to test this hypothesis. Taken together, findings in the NSF, OFT, FST, and food intake under social isolation indicate that stress exposure leads to the expected increases in anxiety-like and depression-like behaviors when *Krtcap3* expression is low.

### KO rats exhibit lower basal CORT and increased CORT response to both chronic stress and an acute psychosocial stress relative to WT rats

Differences in plasma CORT and ACTH between WT and KO rats at euthanasia indicate that *Krtcap3* expression likely plays a role in HPA axis function. As before (Szalanczy, Giorgio et al. 2023), we found that control KO rats had lower basal plasma CORT compared to control WT rats, supported by lower basal ACTH and slightly smaller adrenal glands. At the end of the study, stress-exposed KO rats had greater CORT than control counterparts with no differences in basal ACTH and a mild increase in adrenal gland weight. Contrary to our initial expectations, stress-exposed WT rats did not have greater CORT than control WT rats and in fact had lower basal ACTH than control WT rats, which was surprising as prolonged stress is known to increase ACTH in WKYs (Pardon, Ma et al. 2003, Malkesman, Braw et al. 2006). These findings again support the hypothesis that control WT were as chronically stressed due to the study design. Furthermore, there were also no differences in adrenal gland weight between control and stress-exposed WT rats.

We also confirmed here that KO rats of either stress condition experience a spike in CORT when their cage-mate is removed at euthanasia (Szalanczy, Giorgio et al. 2023), supporting that KO rats are more sensitive to a psychosocial stress. Interestingly, we identified this same pattern in stress-exposed WT rats, but not control WT rats. It is possible that prior stress exposure sensitized the HPA axis to a novel acute stress (Franco, Chen et al. 2016) in WT rats, although the adiposity and behavioral data suggest this is not the case.

It is unclear why control WT rats had such a high basal CORT at euthanasia compared to the other groups. One explanation for this finding is that control WT rats had not fully adjusted to the room change for euthanasia, and the room change stress is responsible for the differences in basal CORT between control WT and KO rats at euthanasia. A second possibility is that the large change in housing environment to a different building coupled with a behavioral test in the final weeks of the study may have more greatly affected control WT rats than KO rats. This also supports that WT and KO rats respond to different stressors differently, as supported by work in other models (Singh, Petrides et al. 1999, Kogler, Muller et al. 2015).

### Nr3c1 expression differs in control and stress-exposed KO rats, with no differences in WT

The pattern of expression for the GR *Nr3c1* may provide insight into the differences seen between WT and KO rats in the current study. We measured *Nr3c1* expression in the pituitary, colon, and liver of WT and KO rats; while we only explored changes in gene expression here, the connection between *Krtcap3* and *Nr3c1* is also supported by evidence of protein-protein interactions elsewhere (Lievens, Van der Heyden et al. 2016).

The pituitary has high *Krtcap3* expression in female WKY rats and is key to the HPA axis. The exact regulation of the GR under chronic stress conditions is unclear. Foundational *in vitro* work demonstrated that GR expression decreases following GC exposure (Sheppard, Roberts et al. 1991, Williams, Franklyn et al. 1991) which aligns with findings in humans that early life stress is associated with methylation of *NR3C1*, leading to decreased gene expression and dysregulated HPA axis function (van der Knaap, Oldehinkel et al. 2015, Alexander, Kirschbaum et al. 2018, Holmes, Shutman et al. 2019, Chatzittofis, Bostrom et al. 2021, Lewis, Breitenstein et al. 2021, Chubar, Vaessen et al. 2023). However, *in vivo* work where animals are subjected to up to a week of stress has shown that expression of the GR increases in the pituitary (Sheppard, Roberts et al. 1990, Nishimura, Makino et al. 2004, Noguchi, Makino et al. 2010). Similarly, in the current study we found that pituitary *Nr3c1* expression increased when KO rats were exposed to stress, with no changes in WT rats. These findings further support that low *Krtcap3* expression may “normalize” the HPA axis of the WKY rat. A change in pituitary *Nr3c1* expression after stress exposure may be related to negative feedback regulation pathway of the HPA axis (Gjerstad, Lightman et al. 2018) and could clarify the role of *Krtcap3*.

*Krtcap3* expression is also high along the gastrointestinal tract of female WKY rats and can also be connected to the stress response (Wiley, Higgins et al. 2016). Other work has shown that chronic stress decreases colonic *Nr3c1* (Zheng, Victor Fon et al. 2017, Muir, Klocke et al. 2023), similar to our findings in the KO rats, lending further credence to the hypothesis that low *Krtcap3* expression “normalizes” the HPA axis of WKY rats. In the colon we found that control KO rats have greater *Nr3c1* expression than control WT rats and that stress exposure decreased *Nr3c1* expression in KO rats to levels comparable to the WT rats. These data may indicate that control KO rats may have improved intestinal integrity over WT rats due to the comparatively high *Nr3c1* expression, but upon exposure to chronic stress and a HFD, *Nr3c1* expression may decrease to minimize damage (Aranda, Arredondo-Amador et al. 2019, Shukla, Meena et al. 2022). The lack of change in *Nr3c1* expression in WT rats after stress exposure supports that WT and KO rats responded to the study design differently. *Krtcap3* may be acting in either the pituitary, the gastrointestinal tract, or both to influence GR expression and the effect of glucocorticoids on metabolism and behavior.

In the current study, we found that chronic stress increased liver *Nr3c1* expression only in the WT rats, where previous work demonstrated increases in both WT and KO rats (Szalanczy, Giorgio et al. 2023). This highlights that the stress administered in this study did not recapitulate the stress present in our previous work and that the role of *Krtcap3* in stress response, and its relationship with *Nr3c1* expression, is tissue-dependent (Costello, Krilis et al. 2022). There remains a need to better understand how different types of stress regulate expression of the GR in different tissues and to identify downstream effects on health.

We did not see changes in *Krtcap3* expression in the pituitary, adrenal, or colon between control and stress-exposed rats. Given that the control and stress-exposed WT rats had similar responses throughout the study, this finding is not surprising. Future work will therefore be needed to determine if stress alters *Krtcap3* expression among these tissues.

## Conclusions

This work confirms that *Krtcap3* expression affects the stress response, with indirect effects on adiposity and behavior. We found here that *Krtcap3*-KO rats were susceptible to the current stress paradigm, with corresponding increases in adiposity and anxiety-like behavior. Despite exposure to the same stressors, there were minimal differences in these measures between control and stress-exposed WT rats, and we hypothesize that this is because control WT rats were also chronically stressed due to study design. This indicates that KO rats may be better able to distinguish between and adapt to stressors than WT rats. That *Nr3c1* expression increases in the pituitary and decreases in the colon in the KO, but not WT rats, supports the hypothesis that low *Krtcap3* expression may “normalize” the HPA axis of WKY rats, although future studies are needed to test this. Importantly, the potential interaction between *Krtcap3* and *Nr3c1*—either at the mRNA level or at protein level—may explain the connection between *Krtcap3* expression, stress, metabolism, and behavior and will be key to further explorations.

## Author Contributions

AMS designed the study, conducted experiments, oversaw assisting researchers, ran statistical analysis, analyzed results, created figures, and wrote the manuscript. LSW designed study, oversaw experimental work, and edited manuscript. MF, TB, SE, and JL assisted with experiments. AB scored the FST, JL and CS scored the NSF test. CD coordinated rat care at Building B and ran the OFT; JLW consulted on behavioral data. MG, JK, and AMG created the rat knock-out model. EER consulted on stress hypotheses. All authors approved final version of the manuscript.

## Supporting information

ARRIVE Guidelines

## Acknowledgements

As always, the authors thank the MCW Genotyping Core for their assistance in genotyping the rats.

---

**Supplementary Figure 1.** *In vivo Krtcap3* knock-out (KO, gray) rats have reduced *Krtcap3* expression compared to wild-type (WT, black) in multiple tissues. Expression was assessed by RT-qPCR in pituitary, adrenal, liver, hypothalamus, ileum, ovary, and retroperitoneal (RetroFat) tissue. Fold change was calculated relative to WT and normalized to pituitary *Krtcap3* expression. KO rats had significantly reduced *Krtcap3* expression in all tissues assessed.

**Supplementary Figure 2.** Adiposity prior to diet start in wild-type (WT, black) and *Krtcap3* knock-out (KO, gray) rats. (A) There were no significant differences between WT and KO rats at wean, nor were there differences in weight between rats assigned to the control group (filled) or stress group (empty). (B) After the first three weeks of stress exposure, prior to high fat diet start, there were still no differences in weight by genotype nor by stress group. (C) There was, however, a main effect of group, where WT and KO rats exposed to stress had increased total fat mass compared to controls. *p < 0.05 represents a main effect of stress.

**Supplementary Figure 3.** Unpredictable chronic mild stress (UCMS) increased weight gain in wild-type (WT, black) and *Krtcap3* knock-out (KO, gray) rats without strong effects on eating behavior, while increasing fat mass in KO rats but not WT. (A) Stress-exposed (empty) rats of both genotypes gained more weight during the UCMS period compared to control (filled) counterparts, against expectations. ****p < 0.0001 represents main effect of stress. (B) Despite the differences in weight gain, stress-exposed rats only demonstrated a mild increase in average weekly food consumption per cage. #p < 0.1 represents main effect of stress. At the EchoMRI analysis at the end of UCMS exposure, (C) there were no differences in total fat mass between WT rats, but stress-exposed KO rats still had greater fat mass than control counterparts. *p < 0.05 represents effect of stress for KO rats. (D) There was a main effect of stress on total lean mass, where stress-exposed rats of both genotypes had greater lean mass compared to control counterparts. *p < 0.05 represents main effect of stress.

