## Supplementary figures and images for "Chronic Stress Increases Adiposity and Anxiety in Rats with Decreased Expression of *Krtcap3*"

### Supplemental Figure 1

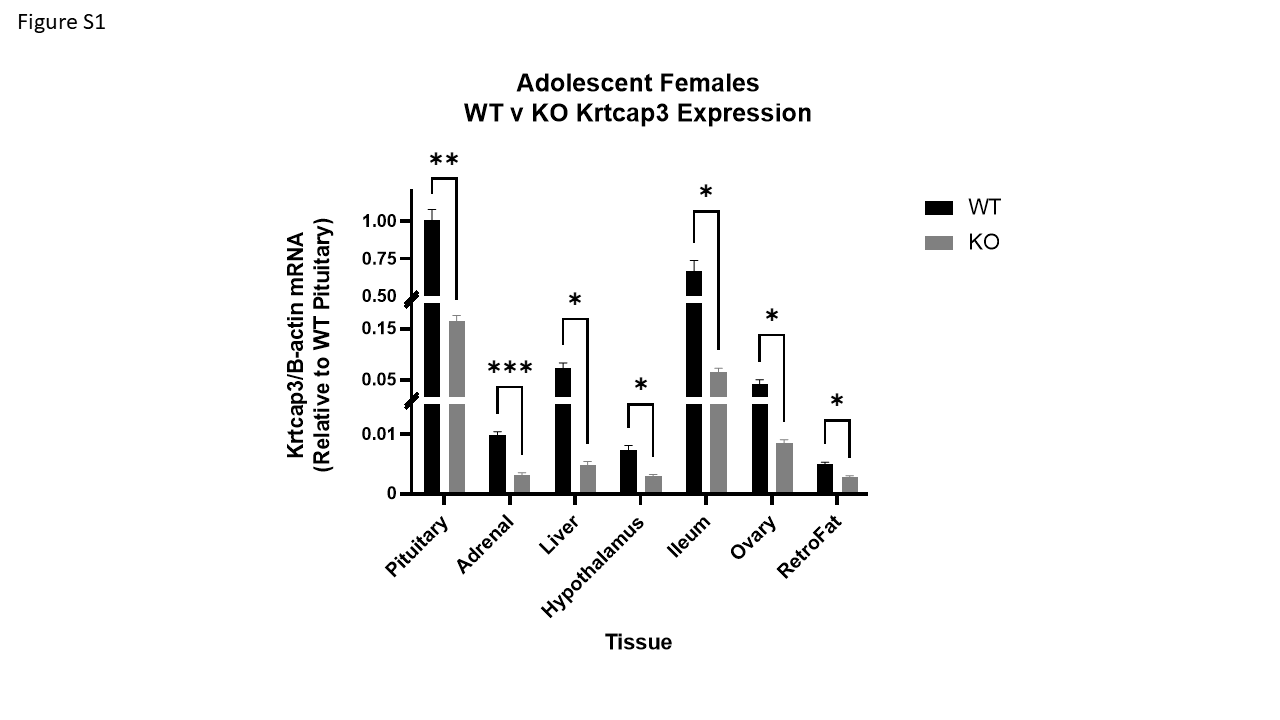

### Supplemental Figure 2

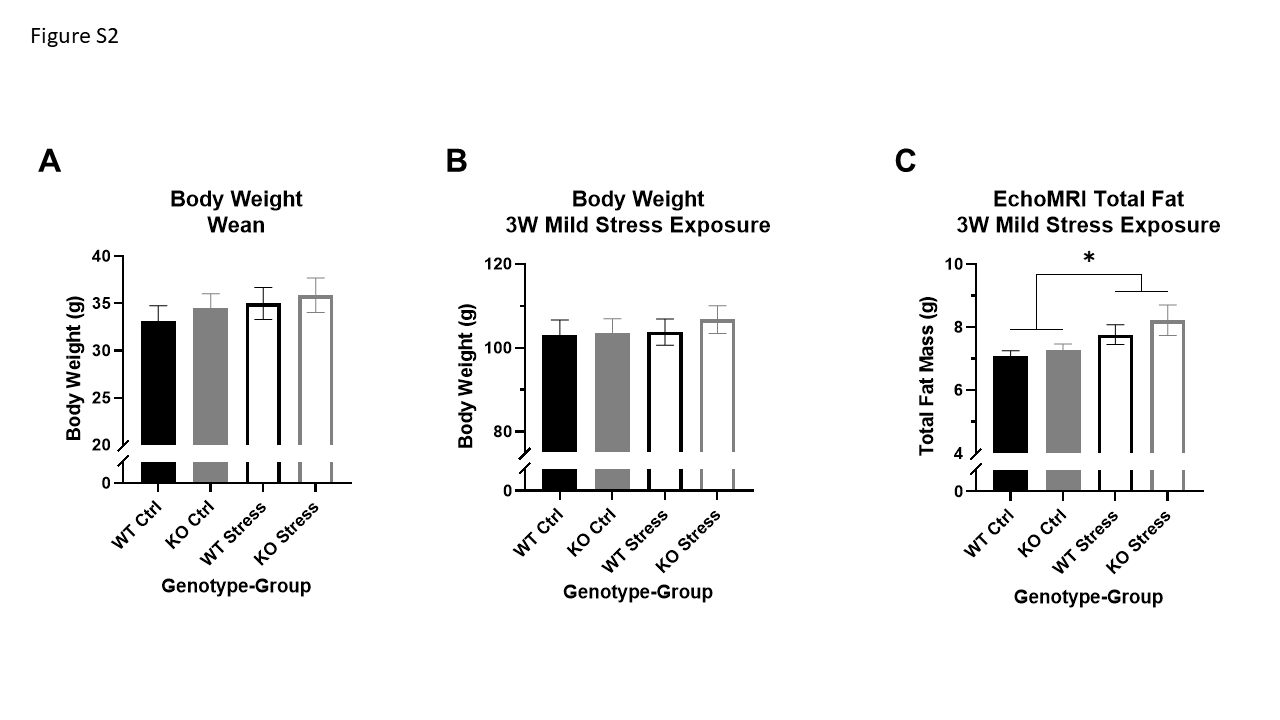

### Supplemental Figure 3

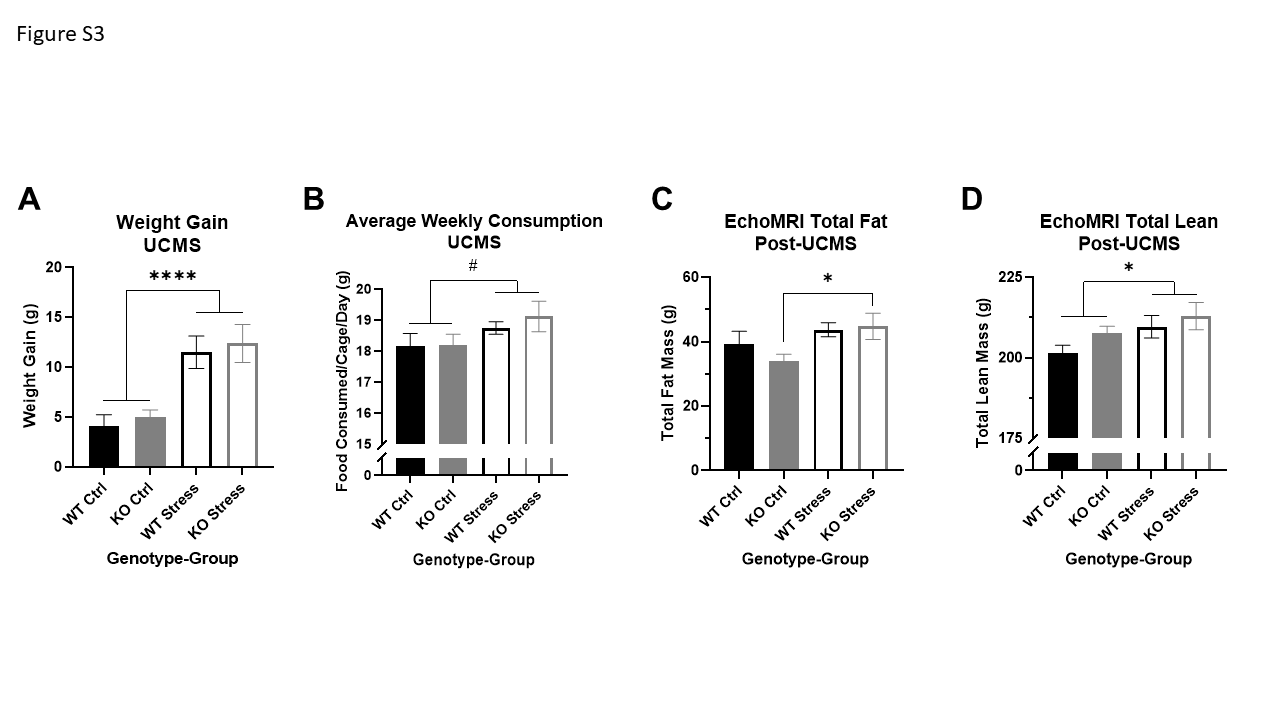
