## Supplementary material for "Chronic Stress Increases Adiposity and Anxiety in Rats with Decreased Expression of *Krtcap3*": ARRIVE Guidelines

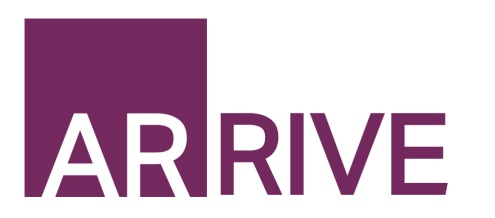

The ARRIVE Guidelines Checklist

Animal Research: Reporting In Vivo Experiments

Carol Kilkenny^1^, William J Browne^2^, Innes C Cuthill^3^, Michael Emerson^4^ and Douglas G Altman^5^

*^1^The National Centre for the Replacement, Refinement and Reduction of Animals in Research, London, UK, ^2^School of Veterinary Science, University of Bristol, Bristol, UK, ^3^School of Biological Sciences, University of Bristol, Bristol, UK, ^4^National Heart and Lung Institute, Imperial College London, UK, ^5^Centre for Statistics in Medicine, University of Oxford, Oxford, UK.*

|  | | ITEM | RECOMMENDATION | Section/ Paragraph |
| --- | --- | --- | --- | --- |
|  | 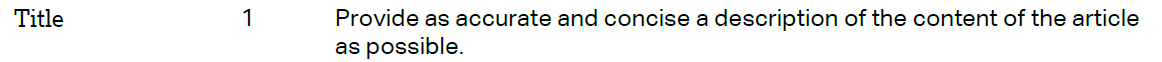 | | | Title Page |
|  | 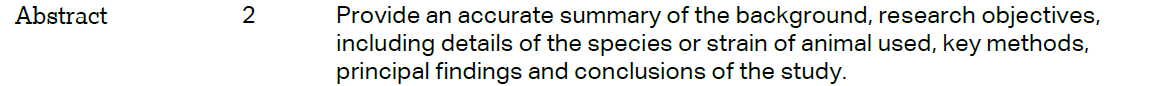 | | | p.3 |
|  | INTRODUCTION | | |  |
|  | 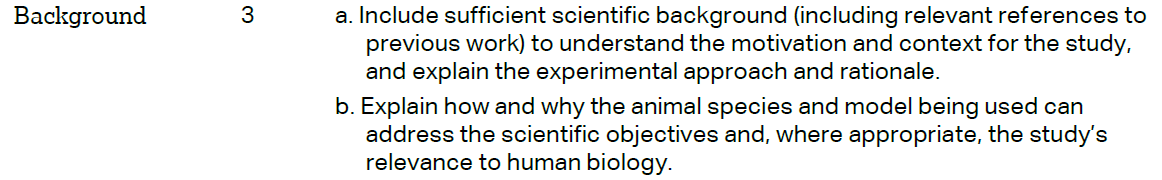 | | | Introduction |
|  | 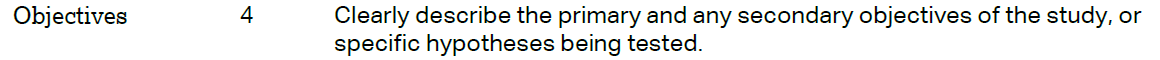 | | | Introduction |
|  | METHODS | | |  |
|  | 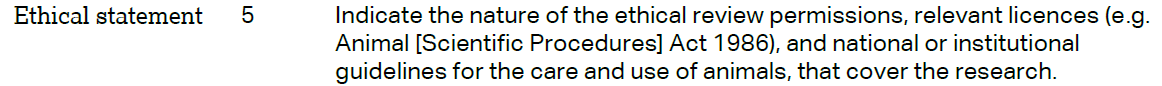 | | | Methods |
|  | 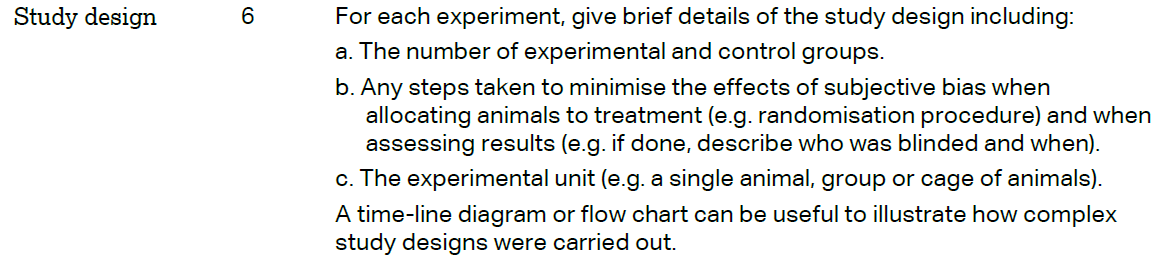 | | | Methods |
|  | 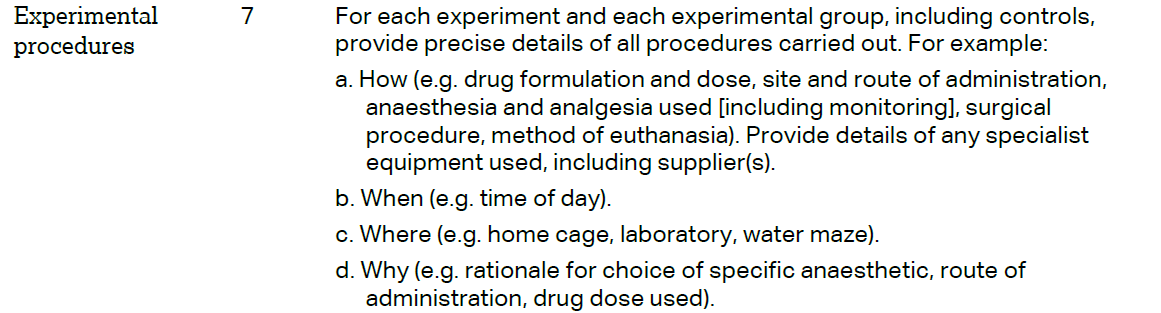 | | | Methods |
|  | 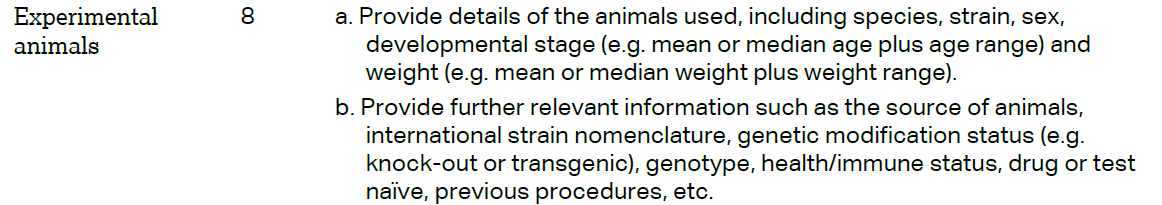 | | | Methods |

The ARRIVE guidelines. Originally published in *PLoS Biology*, June 2010^1^

|  | 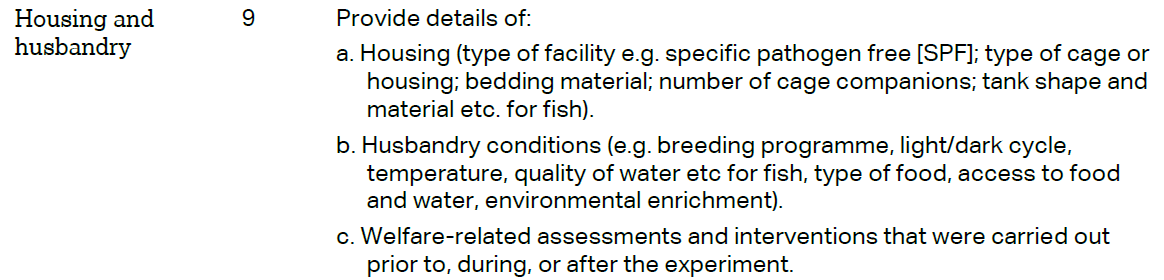 | Methods |
| --- | --- | --- |
|  | 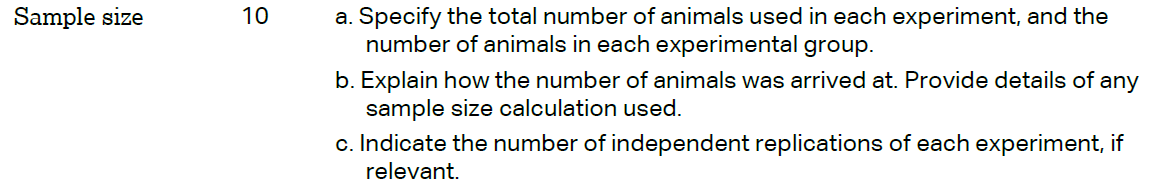 | Methods |
|  | 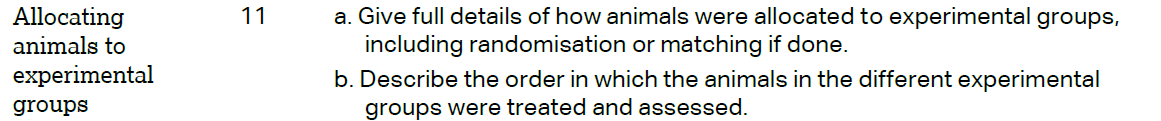 | Methods |
|  | 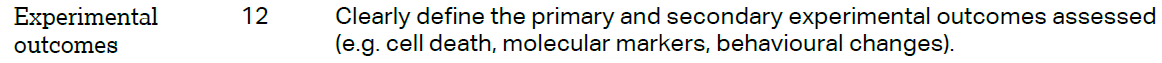 | Methods |
|  | 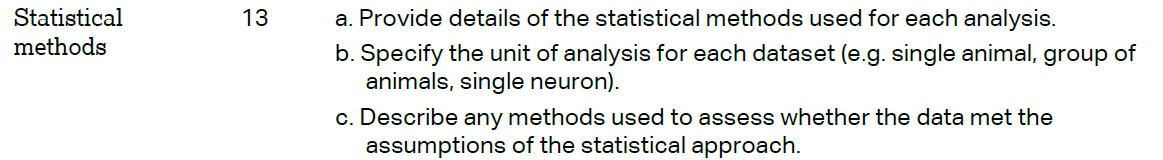 | Methods |
|  | RESULTS |  |
|  | 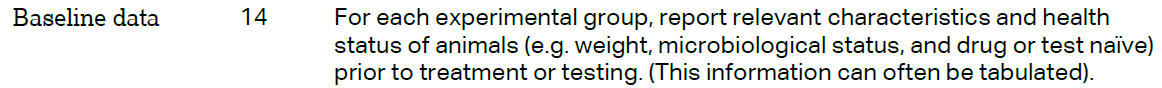 | Results |
|  | 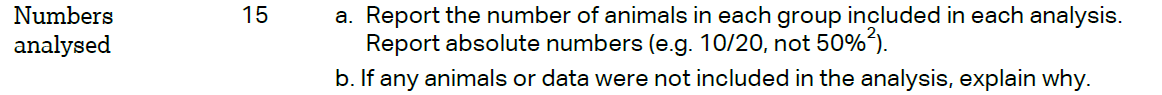 | Results |
|  | 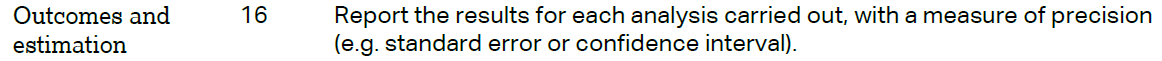 | Results |
|  | 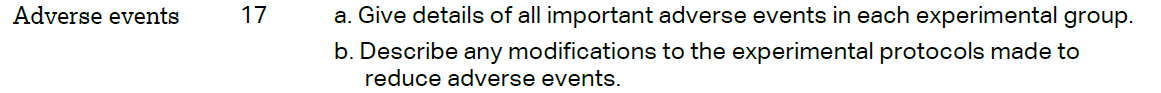 | N/A |
|  | DISCUSSION |  |
|  | 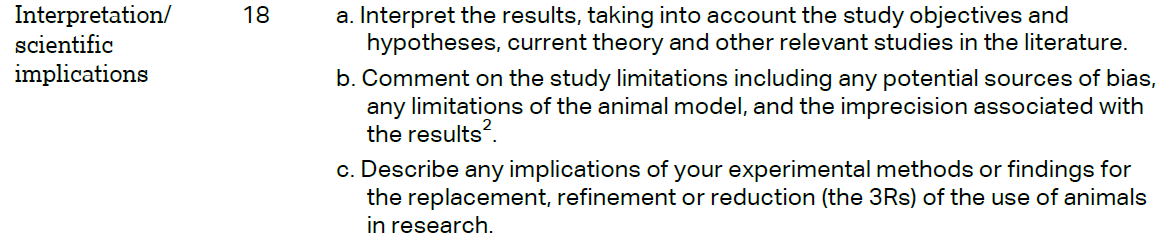 | Discussion |
|  | 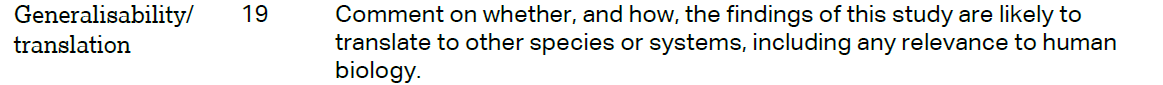 | Discussion |
| 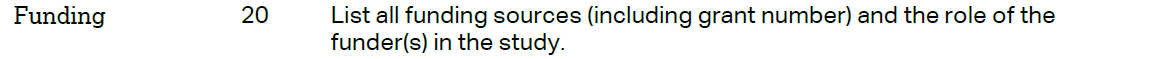 | | Title Page |

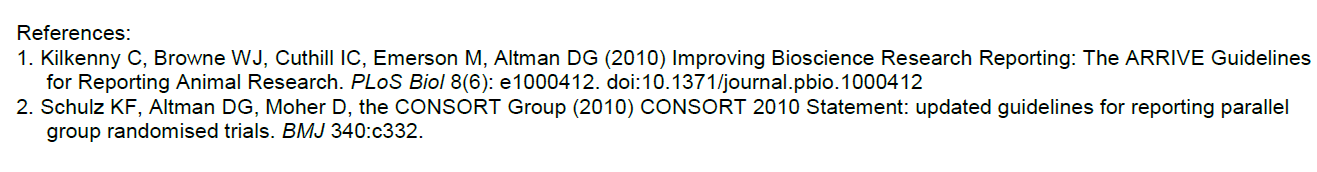

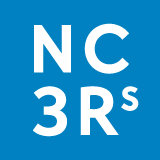
